# Biocatalytic Construction of a CEST MRI Nucleoside Probe: Synthesis and Evaluation of 5-Methyl-5,6-dihydrothymidine

**DOI:** 10.1101/2025.06.25.661137

**Authors:** Aimen Al-Hilfi, E. Alejandro Castellanos Franco, Connor J. Grady, Zinia Mohanta, Michael T. McMahon, Milana Bazayeva, Zhen Li, Kenneth M. Merz, Assaf A. Gilad

**Affiliations:** Department of Chemical Engineering & Material Science, Michigan State University, East Lansing, Michigan 48824, United States; Department of Biomedical Engineering, Michigan State University, East Lansing, Michigan 48824, United States; Russell H. Morgan Department of Radiology and Radiological Science, Johns Hopkins University School of Medicine, Baltimore, Maryland 21205, United States; F.M. Kirby Research Center for Functional Brain Imaging, Kennedy Krieger Research Institute, Baltimore, Maryland 21205, United States; Center for Computational Life Sciences, Lerner Research Institute, The Cleveland Clinic, Cleveland, Ohio 44106, United States; Department of Radiology, Michigan State University, East Lansing, Michigan 48824, United States

## Abstract

Magnetic Resonance Imaging (MRI) is a cornerstone of modern clinical diagnostics, often enhanced by contrast agents. Traditionally, these agents are chemically synthesized, which can involve complex, costly, and environmentally unfriendly processes. Here, we report a novel biocatalytic approach for the efficient, safe, and eco-friendly synthesis of 5-methyl-5,6-dihydrothymidine (5-MDHT), a potent Chemical Exchange Saturation Transfer (CEST) MRI probe for imaging *in vivo* expression of the Herpes Simplex Virus Type-1 Thymidine Kinase (HSV1-TK) reporter gene. We demonstrate that 5-MDHT can be biosynthesized via one- or two-step enzymatic reactions using human purine nucleoside phosphorylase (hPNPase) and the SgvM^VAV^ SAM-dependent methyltransferase. hPNPase catalyzed the base-exchange reaction with catalytic efficiencies *(k*_cat_*/K*_M_) between 138-316 s^−1^ M^−1^, while SgvM^VAV^ methylation of 5,6-dihydrothymidine yielded 5-MDHT with a catalytic efficiency of 26 s^−1^ M^−1^. Molecular dynamics simulations supported the enzymatic binding and selectivity observed experimentally. The resulting 5-MDHT was validated using CEST-MRI, showing a distinct exchangeable imino proton signal at 5.3 ppm. These findings highlight the chemo- and regioselectivity of the biocatalysts and establish biocatalysis as a viable platform for producing clinically relevant MRI contrast agents.

## Introduction

Herpes simplex virus type-1 thymidine kinase (HSV1-TK) is an enzyme that phosphorylates a wide range of nucleoside analogs and has been used as a reporter gene with a variety of radiolabeled nucleotide analogs for different positron emission tomography (PET) applications.^1–3^ However, the complexity of synthesizing radiolabeled probes and their short lifetime limits the accessibility of investigating the use of HSV1-TK as a reporter gene.^4^ Several studies have been constructed to transform the HSV1-TK into an MRI reporter gene by developing a chemical exchange saturation transfer (CEST) agent substrate that can monitor the expression of HSV1-TK.^5–7^ At this point, CEST-MRI has been extensively employed in both preclinical and clinical studies and has become a promising molecular imaging tool that is suitable for imaging enzyme activity, gene expression, and assessing tumor metabolism.^7–12^ CEST-MRI allows the specific detection of metabolites that contain exchangeable (amide, amine, and hydroxyl) protons and the surrounding bulky water protons without the presence of paramagnetic metals.^10,13,14^

5-methyl-5,6-dihydrothymidine (5-MDHT) was identified as an important CEST-MRI contrast agent to monitor the expression and the activity of the HVS1-TK gene.^5,6^ 5-MDHT can be prepared using a four-step chemical synthesis starting from thymidine (**Scheme 1**).^6^ The synthetic steps require protecting the hydroxyl groups of the 2′-deoxyribose moiety of the thymidine by a silyl ether group, the use of the hydrogen gas for the hydrogenation step, and the use of pyrophoric alkyl lithium reagents for the alkylation step (**Scheme 1**).^6^ Despite previous demonstrations in large animals^15^, one of the limitations of 5-MDHT is that the chemical synthesis is hazardous, labor-intensive, and yields low amounts of product.

**Scheme 1:**
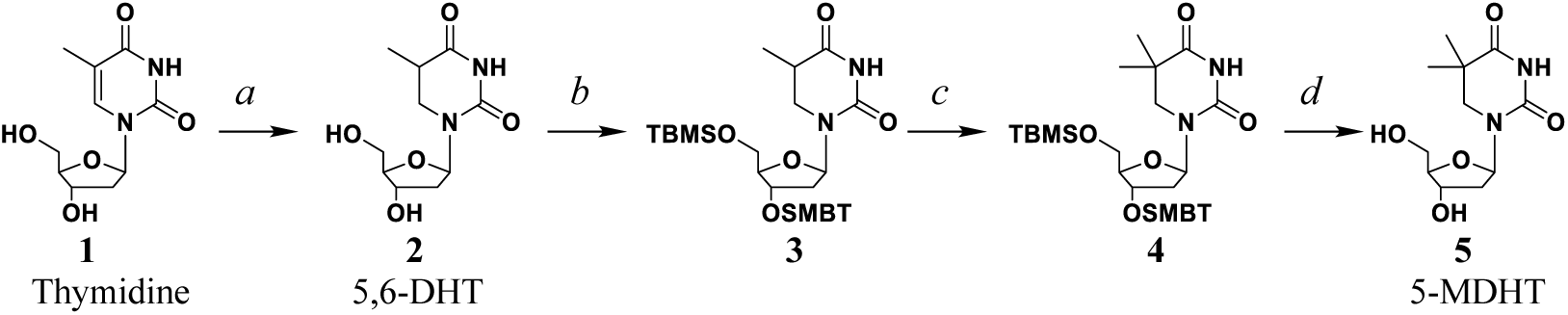
Synthesis of 5-methyl-5,6-dihydrothymidine (5-MDHT). *Reagents and conditions*: (*a*) H_2_ gas, Rhodium on alumina, water/methanol (1:1 v/v), rt, 3 h. (*b*) *tert*-Butyldimethylsilyl chloride (TBMSCl), imidazole, dry pyridine,0 °C to rt over 4 h. (*c*) iodomethane, *sec*-butyllithium (*Sec*-BuLi), dry THF, -78 °C, 7 h. (*d*) triethylamine trihydrofluoride.

Since biocatalysis has become an essential technology for the synthesis of organic compounds with high stereoselectivity and yield^16–22^, we developed a biocatalytic enzymatic approach for 5-MDHT production (**Scheme 2**) and validated it using CEST-MRI. In this study, we developed a two-step catalysis in which we used human dihydropyrimidine dehydrogenase (DPD) to reduce the carbon-carbon double bond between carbon 5 and 6 of the thymidine instead of using hydrogen gas for preparing 5,6-dihydrothymidne (5,6-DHT). We also used human purine nucleoside phosphorylase (hPNPase) base-exchange catalysis between the uracil analog (**7**) and thymidine or 2′-deoxinsosine to produce 5,6-DHT. Then, we used SAM-dependent methyltransferase to transfer the methyl group to the carbon 5 of the 5,6-DHT and produce 5-MDHT (**Scheme 2A**). It occurred to us that we could render the 5-MDHT production more efficient if we made it into a single-step reaction using hPNPase base-exchange catalysis between commercially available uracil analog (**8**) and thymidine or 2′-deoxinsosine (**Scheme 2B**).

**Scheme 2:**
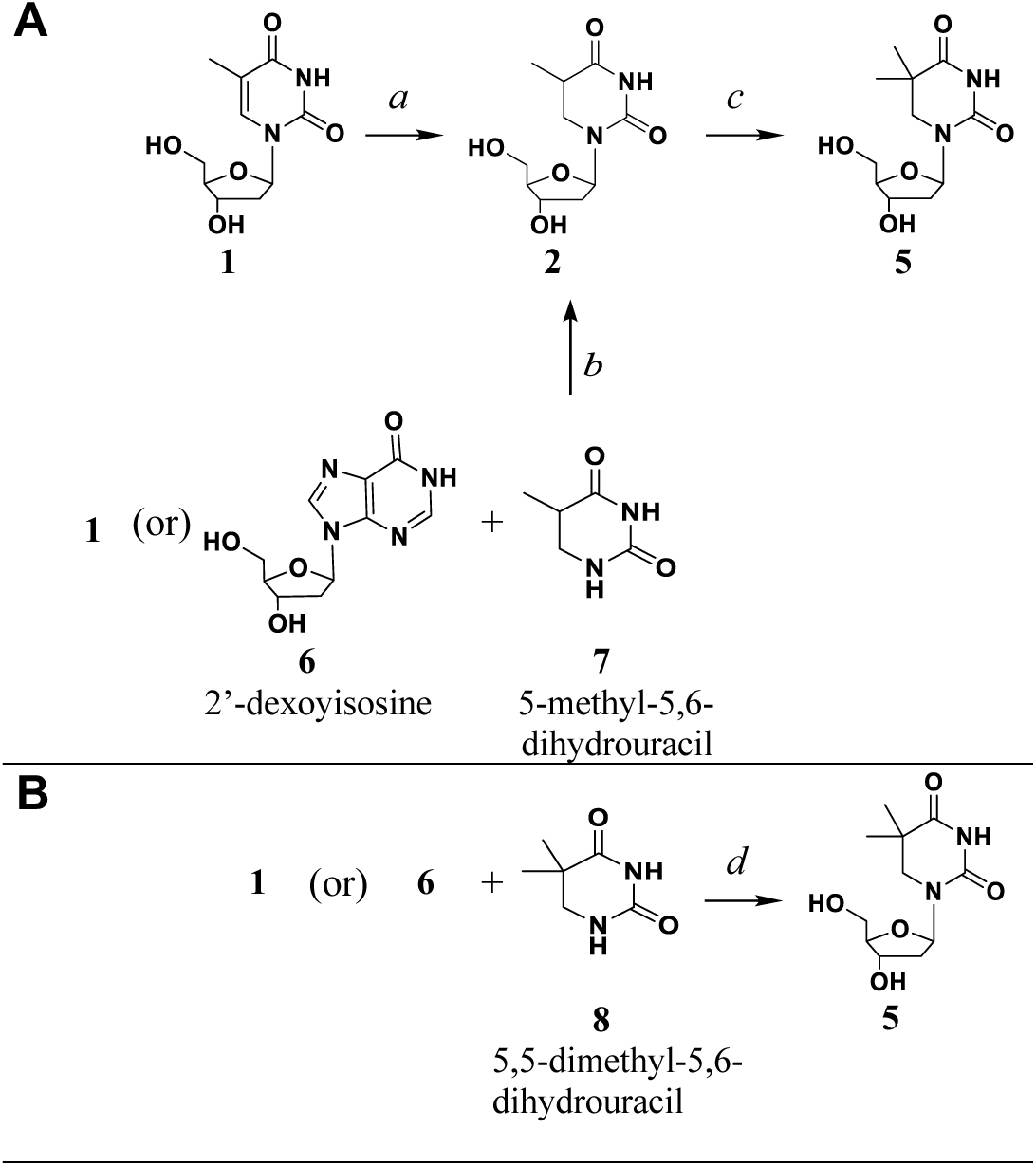
Biocatalysis production methods of 5-MDHT. (A) Proposed two-step catalysis reaction: *Reagents and conditions*: (*a*) Purified hDPD, NADPH, and two flavin cofactors (FAD and FMN), sodium phosphate buffer (pH 7.5), 31 °C, over 16 h; (*b*) Purified hPNPase; sodium phosphate buffer (pH 6.8), 37 °C, over 6 h; (*c*) Purified SgVM, SAM. (B) Proposed single-step catalysis reaction: *Reagents and conditions*: (*d*) Purified hPNPase, sodium phosphate buffer (pH 6.8), 37 °C, over 6 h.

## Experimental Section

### Materials

Thymidine (>99 %) was purchased from Calbiochem (San Diego, CA). 5,6-dihydro-5-methyluracil (>98 %) was purchased from Alfa Aesar (Haverhill, MA). 2′-deoxyinosine (>99 %) was bought from USBiological (Salem, MA). Thymine (>99 %), S-adenosylhomocysteine (>98 %), xanthine oxidase from bovine milk, methyl iodide (>99 %), methanol (>99.9 %), zinc chloride (>98 %), Nicotinamide adenine dinucleotide phosphate tetrasodium salt (NADPH) (>98 %), Flavin adenine dinucleotide disodium salt hydrate (FAD) (>95 %), Riboflavin 5’-monophosphate sodium salt hydrate (FMN) (>70 %), and Iron(II) sulfate heptahydrate (> 99 %) were sourced from Sigma Aldrich (St. Louis, MO). 5,5-Dimethyl-1,3-diazinane-2,4-dione (95 %) was obtained from AmBeed (Arlington Heights, IL). Nickel-affinity chromatography resin (HisPur™ Ni−NTA Resin) was purchased from Thermo Fisher Scientific (Waltham, MA). Isopropyl β-D-1-thiogalactopyranoside (IPTG), kanamycin, ampicillin, Dithiothreitol (DTT), and phenylmethylsulfonyl fluoride (PMSF) were obtained from Gold Bio (St. Louis, MO).

### Expression and purification of the human Dihydro Pyrimidine Dehydrogenase (hDPD)

The pET28a(+)-*hDPD* plasmid containing the hDPD gene was synthesized by GenScript. The hDPD plasmid was transformed into *E*. *coli* BL21 (DE3) cells (Thermo Scientific), plated onto LB agar containing (50 μg/mL) kanamycin and grown for 16 h with shaking at 37 °C. Single colonies were then used to inoculate (4×30 mL) of LB media with (50 μg/mL) kanamycin and cultures were incubated for overnight with shaking at 37 °C. The inoculum cultures were added to fresh LB media (4×1 L) containing kanamycin (50 μg/mL). The cells were incubated at 37 °C until OD_600_ ≈0.6, IPTG (250 μM final concentration) was added, and the strains were incubated at 16 °C for 16 h. The cultures were centrifuged (4,000 g) for 30 min at 4 °C to pellet the cells. The cells were resuspended in 100 mL of lysis buffer [50 mM) sodium phosphate (pH 8.0), (500 mM) NaCl, (10 mM) imidazole, and 5 % (v/v) glycerol], and lysed by sonication (Misonix Sonicator (Danbury, CT): 10 s on, 20 s rest for 30 cycles) on ice. The cell debris was removed by centrifugation (15,000 *g*) for 45 min at 4 °C. The supernatant was loaded onto a column containing nickel−nitrilotriacetic acid (Ni−NTA) resin (3 mL) and eluted by gravity flow. The column was washed twice with 50 mL of Wash 1 Buffer [(500 mM) NaCl, (50 mM) sodium phosphate (pH 8.0), (10 mM) imidazole, and 5 % (v/v) glycerol] and 25 mL of Wash 2 Buffer [(500 mM) NaCl, (50 mM) sodium phosphate (pH 8.0), (50 mM) imidazole, and 5 % (v/v) glycerol]. The protein was eluted with Elution Buffer [500 mM) NaCl, (50 mM) sodium phosphate (pH 7.5), (250 mM) imidazole, and 5 % (v/v) glycerol]. Fractions containing enzymes of a molecular weight consistent with that of hDPD (∼112 kDa) were combined, concentrated, and desalted using Amicon Ultra-15 10K filters. The concentrated sample was injected into a pre-calibrated Superdex 200 pg column (ÄKTA Pure) with (50 mM) sodium phosphate (pH 7.5) buffer. Fractions containing hDPD were pooled and concentrated. The quantity of hDPD (∼20 mg) was measured using a NanoDrop spectrophotometer, and the purity of the enzyme was further confirmed by SDS-PAGE analysis (**Figure S1** of the Supporting Information). These procedures were repeated as needed to obtain sufficient catalysts for downstream kinetic analyses and scale-up procedures.

### Testing hDPD activity with thymidine

The hDPD reduction catalysis activity was tested by incubating thymidine (400 μM) with (50 μg/mL) purified hDPD, NADPH (400 μM), DTT (1mM), FAD (50 μM), FMN (50 μM), and FeSO_4_ (200 μM) in 3 mL of assay buffer [(50 mM) NaH_2_PO_4_/Na_2_HPO_4_ buffer (pH 7.5)]. The assay was mixed and incubated at 31 °C on a rocking shaker for 16 h. The reaction stopped with methanol (3×1 mL); the organic fractions were combined, and the solvent was removed under a N_2_ gas stream. The remaining residue was resuspended in 100 μL of DI-water, and an aliquot was analyzed by liquid chromatography-electrospray ionization mass spectrometry (LC-ESI-MS) to monitor the production of a mass consistent with 5,6-dihydrothymidine (5,6-DHT) (**Figure S2** of the Supporting Information).

### Expression and purification of the human Purine Nucleoside Phosphorylases (hPNPase)

The pCRT7/NT-TOPO-*hPNPase* plasmid (plasmid # 64076, GenBank ID: AH001522.1) containing the *hPNPase* gene was purchased from Addgene. The hPNPase plasmid was transformed into *E. coli* BL21 (DE3) cell, plated onto LB agar containing ampicillin (100 μg/mL) and grown at 37 °C for 16 h. Single colonies were used to inoculate (6×25 mL) of LB media containing ampicillin (100 μg/mL), and the cultures were incubated overnight with shaking at 37 °C. The overnight cultures were used to re-inoculate fresh LB media (6×1 L) containing ampicillin (100 μg/mL). The cells were incubated at 37 °C shaker and allowed to reach an OD_600_ of 0.6. IPTG (100 μM final concentration) was added, and the strains were incubated at 16 °C for 16 h. Cells were harvested by centrifugation (4,000 *g*) for 30 min at 4 °C. The cell pellets were resuspended in 100 mL of lysis buffer [50 mM) sodium phosphate (pH 8.0), (500 mM) NaCl, (10 mM) imidazole, and 5 % (v/v) glycerol], and lysed by sonication (Misonix Sonicator (Danbury, CT)): 10 s on, 20 s rest for 30 cycles) on ice. The cell debris was removed by centrifugation (15,000 *g*) for 45 min at 4 °C. The supernatant was loaded onto a column containing nickel−nitrilotriacetic acid (Ni−NTA) resin (3 mL) and eluted by gravity flow. The column was washed twice with 50 mL of Wash 1 Buffer [(500 mM) NaCl, (50 mM) sodium phosphate (pH 8.0), (10 mM) imidazole, and 5 % (v/v) glycerol] and 25 mL of Wash 2 Buffer [(500 mM) NaCl, (50 mM) sodium phosphate (pH 8.0), (50 mM) imidazole, and 5 % (v/v) glycerol]. The protein was eluted with Elution Buffer [500 mM) NaCl, (50 mM) sodium phosphate (pH 7.5), (250 mM) imidazole, and 5 % (v/v) glycerol]. Fractions containing enzymes of a molecular weight consistent with that of hPNPase (∼33 kDa) were combined, concentrated, and desalted using Amicon Ultra-15 10K filters. The concentrated sample was injected into a pre-calibrated Superdex 200 pg column (ÄKTA Pure) with (50 mM) sodium phosphate (pH 7.5) buffer. Fractions containing hPNPase were pooled and concentrated. The quantity of hPNPase (∼32 mg) was measured using a NanoDrop spectrophotometer, and the purity of the enzyme was further confirmed by SDS-PAGE analysis (**Figure S1** of the Supporting Information). These procedures were repeated as needed to obtain sufficient catalysts for downstream kinetic analyses and scale-up procedures.

### Screening hPNPase activity with uracil analogs (7 and 8)

The hPNPase catalysis activity with uracil analogs was tested by incubating (2 mM) of **7** and **8** separately with (70 μg/mL) purified hPNPase, and [(2 mM) 2′-deoxyinosine and (0.34 mM) xanthine oxidase] or (2 mM) thymidine in 3 mL of assay buffer [(50 mM) NaH_2_PO_4_/Na_2_HPO_4_ (pH 6.8)]. The assays were mixed and incubated at 37 °C on a rocking shaker for six hours. The reactions were stopped with methanol (3×1 mL); the organic fractions were combined, and the solvent was removed under a N_2_ gas stream. The remaining residue was resuspended in 100 μL of DI-water, and an aliquot was analyzed by liquid chromatography-electrospray ionization mass spectrometry (LC-ESI-MS) to monitor the production of a mass consistent with 5,6-dihydrothymidine (5,6-DHT) and 5-methyl-5,6-dihydrothymidine (5-MDHT) (**Figures 1**).

**Figure 1:**
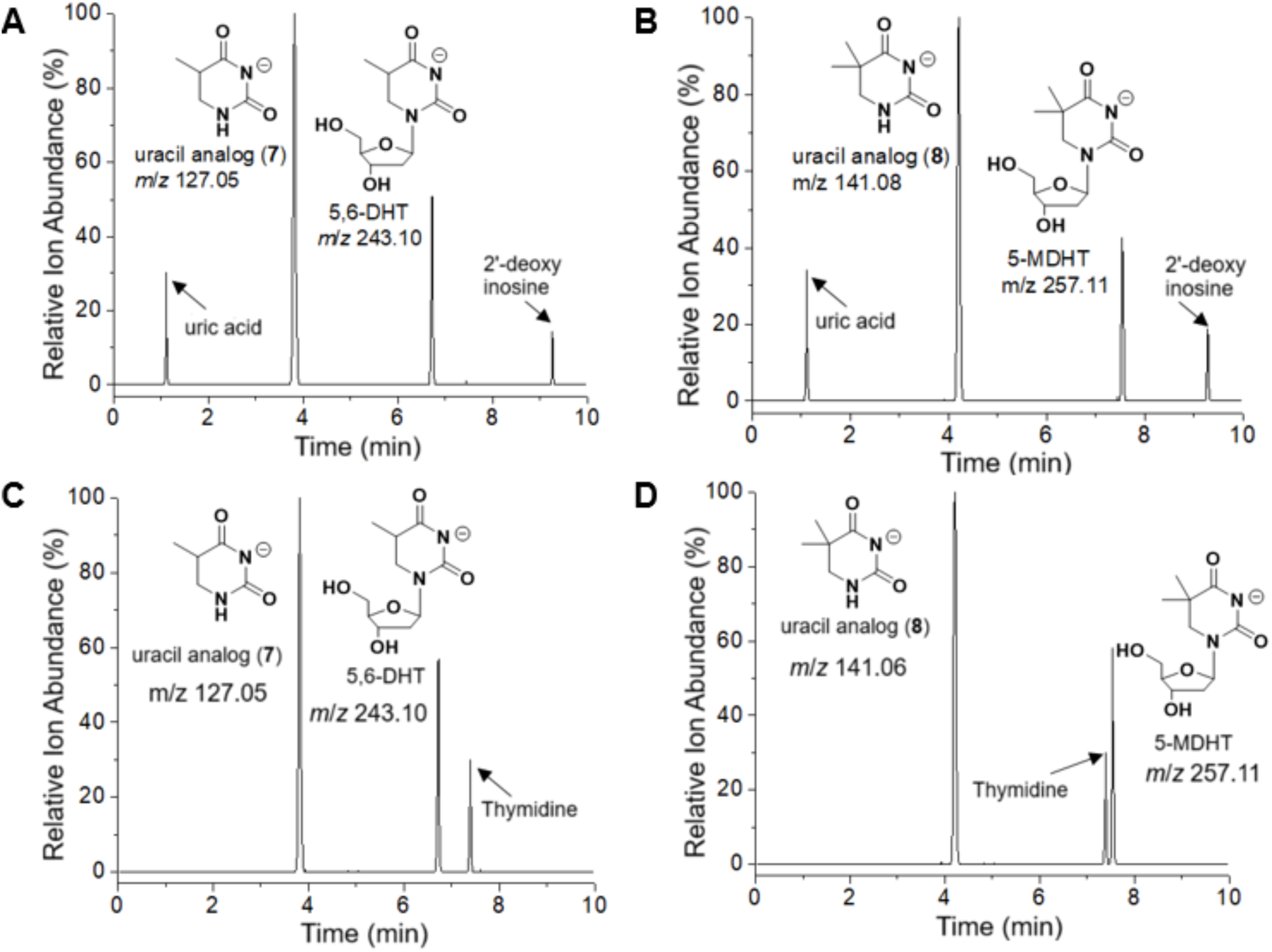
LC/ESI-MS (selected ion mode for *m/z* [M - H]^-^) of the hPNPase base-exchange catalysis (A) between 2′-deoxyinosine and 5,6-dihydro-5-methyluracil to make 5,6-DHT, or (B) 5,5-dimethyl-1,3-diazinane-2,4-dione to make 5-MDHT; (C) between thymidine and 5,6-dihydro-5-methyluracil to make 5,6-DHT, or (D) between thymidine and 5,5-dimethyl-1,3-diazinane-2,4-dione to make 5-MDHT.

### Kinetic evaluation of hPNPase with uracil analogs (7 and 8)

The steady-state conditions for protein concentration and time were established for hPNPase and uracil analogs (**7** and **8**) separately incubated at low (0.05 mM) and high (1 mM) concentrations in 10 mL of assay buffer [(50 mM) NaH_2_PO_4_/Na_2_HPO_4_ (pH 6.8)] containing hPNPase (350 μg/mL), and [2′-deoxyinosine (1 mM) and xanthine oxidase (0.17 mM)] or (2 mM) thymidine at 37 °C on a rocking shaker. Aliquots (1 mL) from each assay were collected, and the biosynthetic reaction was quenched with 500 μL of methanol at 10, 15, 30, and 45 min and 1, 2, 3, 5, 7, and 10 h. Then, each sample was extracted with methanol (3×1 mL). The organic fractions were combined, and the solvent was removed under an N_2_ gas stream. The resultant residue from each assay was separately resuspended in DI-water (100 μL) and quantified by LC-ESI-MS.

A stop time was established for the steady-state time range by incubating hPNPase (350 μg/mL) and [(1 mM) 2′-deoxyinosine and (0.17 mM) xanthine oxidase] or (1 mM) thymidine with varying concentrations of uracil analogs (**7** and **8**) (0.05–1 mM), respectively, in triplicate assays at 37 °C on a rocking shaker for four h. As described above, assay products were extracted from the reaction mixture and quantified by LC-ESI-MS (**Figure S3-S6** of the Supporting Information). The kinetic parameters (*K*_M_ and *k*_cat_) were calculated by nonlinear regression with Origin Pro 9.0 software (Northampton, MA) using the Michaelis-Menten equation: *v*_0_ = *k_cat_*[*E*_0_] [*S*]/(*K_M_* + [*S*]).

### Scale up the hPNPase catalysis production of 5,6-DHT and 5-MDHT

A large-scale preparative hPNPase enzymatic assay was performed under the following conditions: (2 mM) of uracil analogs (**7** and **8**) were incubated separately in 100 mL (25 test tubes, a 4 mL assay in each tube) of assay buffer [(50 mM) NaH_2_PO_4_/Na_2_HPO_4_ (pH 6.8)] containing hPNPase (520 μg/mL, ∼52 mg total), [(2 mM) 2′-deoxyinosine, and (0.34 mM) xanthine oxidase] or (2 mM) thymidine at 37 °C on a rocking shaker for four hours. The reaction stopped with methanol (2×4 mL). The organic fractions were combined, and the solvent was removed under a stream of N_2_ gas. The residue was purified by C18 column (RediSep Gold®, C18 Aq, 20-40 μm, 100 Å) flash chromatography (Combi flash Nextgen 300+) to yield ≥95 % pure product as determined by NMR (**Figures S8-S13** of the Supporting Information). The purified residue was dissolved in DI-water (100 μL), and an aliquot was analyzed by LC-ESI-MS/MS for fragmentation analysis and monoisotopic mass calculation (**Figure S16-S17** of the Supporting Information).

### Expression and Purification of the SAM-dependent methyltransferase from *Streptomyces griseoviridis* (SgvM^VAV^) and thiopurine S-methyltransferase from *Pseudomonas aeruginosa* (*Pa*HMT)

The primers encoding SgvM^VAV^ (GenBank ID: AGN74875.1) and *Pa*HMT (GenBank ID: WP_003114764.1) genes were ordered from IDT and separately cloned into pET-28a(+) via Gibson Assembly. Plasmids containing the SgvM^VAV^ and *Pa*HMT were separately transformed into *E*. *coli* BL21 (DE3) cells, plated onto LB agar containing (50 μg/mL) kanamycin and grown for 16 h with shaking at 37 °C. Single colonies were then used to inoculate (5×30 mL) of LB media with (50 μg/mL) kanamycin and cultures were incubated for overnight with shaking at 37 °C. The inoculum cultures were added to fresh LB media (5×1 L) containing kanamycin (50 μg/mL). The cells were incubated at 37 °C until OD_600_ ≈0.6, IPTG (250 μM final concentration) was added, and the strains were incubated at 16 °C for 16 h. The cultures were centrifuged (4,000 g) for 30 min at 4 °C to pellet the cells. The cells were resuspended in 100 mL of lysis buffer [50 mM) sodium phosphate (pH 8.0), (500 mM) NaCl, (10 mM) imidazole, and 5 % (v/v) glycerol], and lysed by sonication (Misonix Sonicator (Danbury, CT): 10 s on, 20 s rest for 30 cycles) on ice. The cell debris was removed by centrifugation (15,000 *g*) for 45 min at 4 °C. The supernatant was loaded onto a column containing nickel−nitrilotriacetic acid (Ni−NTA) resin (3 mL) and eluted by gravity flow. The column was washed twice with 50 mL of Wash 1 Buffer [(500 mM) NaCl, (50 mM) sodium phosphate (pH 8.0), (10 mM) imidazole, and 5 % (v/v) glycerol] and 25 mL of Wash 2 Buffer [(500 mM) NaCl, (50 mM) sodium phosphate (pH 8.0), (50 mM) imidazole, and 5 % (v/v) glycerol]. The protein was eluted with Elution Buffer [500 mM) NaCl, (50 mM) sodium phosphate (pH 8.0), (250 mM) imidazole, and 5 % (v/v) glycerol]. Fractions containing enzymes of a molecular weight consistent with that of SgvM^VAV^ (∼39 kDa) and *Pa*HMT (∼28 kDa) were combined, concentrated, and desalted using Amicon Ultra-15 10K filters. The concentrated sample was injected into a pre-calibrated Superdex 200 pg column (ÄKTA Pure) with (50 mM) sodium phosphate (pH 8.0) buffer. Fractions containing SgvM^VAV^ and *Pa*HMT were pooled and concentrated. The quantity of SgvM^VAV^ (∼25 mg) and *Pa*HMT (∼28 mg) were measured using a NanoDrop spectrophotometer, and the purity of the enzymes was further confirmed by SDS-PAGE analysis (**Figure S1** of the Supporting Information). These procedures were repeated as needed to obtain sufficient catalysts for downstream kinetic analyses and scale-up procedures.

### Evaluation of SgvM^VAV^ and *Pa*HMT with 5,6-DHT

A solution of 5,6-DHT (2 mM), purified SgvM^VAV^ (100 μg/mL), purified *Pa*HMT (100 μg/mL), (25 μM) S-adenosylhomocysteine, and (5 mM) methyl iodide in 2 mL of assay buffer [(50 mM) NaH_2_PO_4_/Na_2_HPO_4_ (pH 8.0)] was incubated separately with ZnCl_2_ (2 mg), and without ZnCl_2_. The assays were mixed at 31 °C on a rocking shaker for eight hours. The reaction stopped with methanol (4×1 mL); the organic fractions were combined, and the solvent was removed under an N_2_ gas stream. The remaining residue was resuspended in 100 μL of DI-water, and an aliquot was analyzed by liquid chromatography-electrospray ionization mass spectrometry (LC-ESI-MS) to check if the SgvM^VAV^ and *Pa*HMT catalyzed the formation of 5-merthyl-5,6-dihydrothymidine (5-MDHT).

### Kinetic evaluation of SgvM^VAV^ and *Pa*HMT with 5,6-DHT

To establish the steady-state conditions for protein concentration and time for SgvM^VAV^ and *Pa*HMT, the 5,6-DHT was separately incubated at low (0.05 mM) and high (1 mM) concentrations in 10 mL of assay buffer [50 mM NaH_2_PO_4_/Na_2_HPO_4_ (pH 8.0)] containing purified SgvM^VAV^ (150 μg/mL), purified *Pa*HMT (150 μg/mL), (25 μM) S-adenosylhomocysteine, (5 mM) methyl iodide, and ZnCl_2_ (10 mg) at 31 °C on a rocking shaker. Aliquots (1 mL) from each assay were collected, and the biosynthetic reaction was quenched with 500 μL of methanol at 10, 15, 30, and 45 min and 1, 2, 3, 5, 7, and 10 h. Then, each sample was extracted with methanol (3×1 mL). The organic fractions were combined, and the solvent was removed under a N_2_ gas stream. The resultant residue from each assay was separately resuspended in DI-water (100 μL) and quantified by LC-ESI-MS.

A stop time was established for the steady-state time range. SgvM^VAV^ (150 μg/mL), *Pa*HMT (150 μg/mL), (25 μM) S-adenosylhomocysteine, (5 mM) methyl iodide, and ZnCl_2_ (10 mg) were incubated with varying concentrations of 5,6-DHT (0.05–1 mM), respectively, in triplicate assays at 31 °C on a rocking shaker for four hours. As described above, assay products were extracted from the reaction mixture and quantified by LC-ESI-MS (**Figure S7** of the Supporting Information). The kinetic parameters (*K*_M_ and *k*_cat_) were calculated by nonlinear regression with Origin Pro 9.0 software (Northampton, MA) using the Michaelis-Menten equation: *v*_0_ = *k_cat_*[*E*_0_] [*S*]/(*K_M_* + [*S*])

### Scale-up the SgvM^VAV^ and *Pa*HMT catalytic production of 5-MDHT

A concentrated solution of purified SgvM^VAV^ (750 μg/mL, ∼75 mg total) and purified *Pa*HMT (750 μg/mL, ∼75 mg total) were incubated in 100 mL (25 test tubes, a 4 mL assay in each tube) of assay buffer [(50 mM) NaH_2_PO_4_/Na_2_HPO_4_ (pH 8.0)] containing (2 mM) 5,6-DHT, (25 μM) S-adenosylhomocysteine (SAH), and (5 mM) methyl iodide and ZnCl_2_ (100 mg) at 31 °C on a rocking shaker for 8 h. The reactions were stopped with methanol (2×3 mL). The organic fractions were combined, and the solvent was removed under a stream of N_2_ gas. The residue was purified by C18 column (RediSep Gold®, C18 Aq, 20-40 μm, 100 Å) flash chromatography (Combi flash Nextgen 300+) to yield ≥94 % pure product as determined by NMR (**Figures S14-S15** of the Supporting Information). The purified residue was dissolved in DI-water (100 μL), and an aliquot was analyzed by LC-ESI-MS/MS for fragmentation analysis and monoisotopic mass calculation (**Figure S17** of the Supporting Information).

### Molecular dynamic simulation studies

Structure optimization on 5,6-dihydrothymidine, 5,5-dimethyl-5,6-dihydrouracil, and 2′-deoxyribose-1α-phosphate were conducted using Gaussian 16 in a four-step pattern^23^, starting from HF 3-21G* single point to HF 3-21G* optimization, then to B3LYP 3-21G*, and finally to B3LYP 6-31G*.^24,25^

Molecular dynamics (MD) simulations were performed using AMBER24. The topology and coordinate files for hPNPase (PDB ID: 1RFG) and SgvM^VAV^ (PDB ID: 8FTV) were constructed using the tLEaP program in the AMBER24 package. The ff19SB forcefield and OPC water model^26^ were used. The system was prepared in three steps. First, the antechamber, prepin, and parkmchk2 programs in the AmberTools23 package^27^ were used to generate the charge and force constants for all the reactants using AM1BCC charge model and GAFF2 forcefield. Minimization was done in five stages, gradually removing restrictions from the protein backbone to the side chain. Each step yields 10,000 steps of the steepest descendent and 10,000 steps of conjugate gradient methods. A quick 9-ps *NPT* simulation was conducted to avoid the formation of bubbles during heating. Afterward, a 360-ns *NVT* heating was performed with the temperature increasing gradually from 0 to 300 K. Then another 20-ns simulation was performed to equilibrate the system in the *NPT* ensemble, and the last 2,000 frames were used for distance analysis. The PME method and PBC were used for the simulations, and the Langevin algorithm with a 2.0 ps^-1^ friction frequency coefficient was used to maintain the temperature.^28^ The Berendsen barostat method was used for pressure control with a relaxation time of 1.0 ps.^29^ The time step was 1.0 fs, with the SHAKE function constraining the hydrogen atom bonds.^30^

### Evaluating 5-MDHT as a CEST-MRI contrast agent

The 5-MDHT produced by two and single-step biocatalysis reactions was dissolved at a concentration of 50 mM in phosphate-buffered saline (PBS) (pH 7.4) and placed in 3-mm NMR tubes for performing CEST MRI experiments on a vertical 11.7 T scanner (Bruker Avance system) at 37 °C as previously described^10^ with TR/TE set to 5000/17.27 ms. The saturation module employed a block pulse with peak saturation power (*B_1_*) = 4 µT, saturation time (tsat) = 3 s, and the saturation offset incremented from 4000 to -4000 Hz from water, with a step size of 100 Hz. Data processing was performed using custom-written scripts in MATLAB (Mathworks) as described previously (add references 18 in MM suggestion). Mean CEST spectra were plotted using a ROI drawn on each tube after *B*_0_ correction. MTR_asym_ = 100 × (*S*^−Δ*ω*^ − *S*^Δ*ω*^)/*S*_0_ was computed at different offsets Δ*ω*.

## Results and Discussion

### Production of 5,6-dihydrothymidine from thymidine by hDPD

The reduction catalysis of hDPD was tested by incubating thymidine and NADPH with hDPD, FAD, FMN, DTT, and FeSO_4_. The bio-catalyzed product was screened by LC-ESI-MS selected ion monitoring. Selected ion *m/z* 243.10 was identified in the LC-ESI-MS profile and assigned to the [M-H]^-1^ ion for the 5,6-dihydrothymidine (5,6-DHT), which shows that hDPD can catalyze the reduction of the thymidine to the 5,6-DHT (**Figure S2** of the Supporting Information). However, the presence of dioxygen limited the hDPD reduction activity. This is because the iron-sulfur cluster (electron bridge between FAD and FMN) is unstable in the presence of oxygen, and iron-sulfur protein purification must be done under anaerobic conditions as described in previous studies.^31,32^ Therefore, these limitations will decrease the hDPD reduction productivity as the LC-ESI-MS results showed a low-intensity signal of the production of the 5,6-DHT (**Figure S2** of the Supporting Information).

### Screening and kinetics evaluation of hPNPase activity with uracil analogs (7 and 8)

#### Activity

The hPNPase ability to transfer 2’-deoxyribose to the uracil base substrates was tested by incubating separately uracil analogs (**7** and **8**) with hPNPase, and [2′-deoxyinosine/xanthine oxidase] or thymidine. LC-ESI-MS analysis was used to assess the relative turnover of substrate to the product. Selected ions *m/z* 243.11 and *m/z* 257.11 were identified in the LC-ESI-MS profiles and putatively assigned to the [M-H]^-1^ for the 5,6-DHT and 5-MDHT respectively (**Figure 1**).

#### Kinetics

The *K*_M_ and *k*_cat_ of the hPNPase catalysis reaction were calculated under steady-state conditions by incubating hPNPase with varying concentrations of uracil analogs (**7** and **8**) and [2′-deoxyinosine/xanthine oxidase] or thymidine in separate assays (**Figure S3-S6** of Supporting Information). The catalytic efficiency (*k*_cat_/*K*_M_) values of the hPNPase catalyzed base-exchange reaction between uracil analogs and 2′-deoxyinosine were higher than those values of base exchange with thymidine (**Table 1**). This is because the hPNPase catalyzed base exchange with 2′-deoxyinosine produces hypoxanthine as a by-product, which is converted into uric acid by adding xanthine oxidase (**Scheme 3A**). As a result, that shifts the equilibrium reaction further to the right and increases the turnover number of the bio-catalyzed products. On the other hand, in the hPNPase base-exchange catalysis with thymidine, thymidine is the by-product (**Scheme 3B**).

**Table 1:**
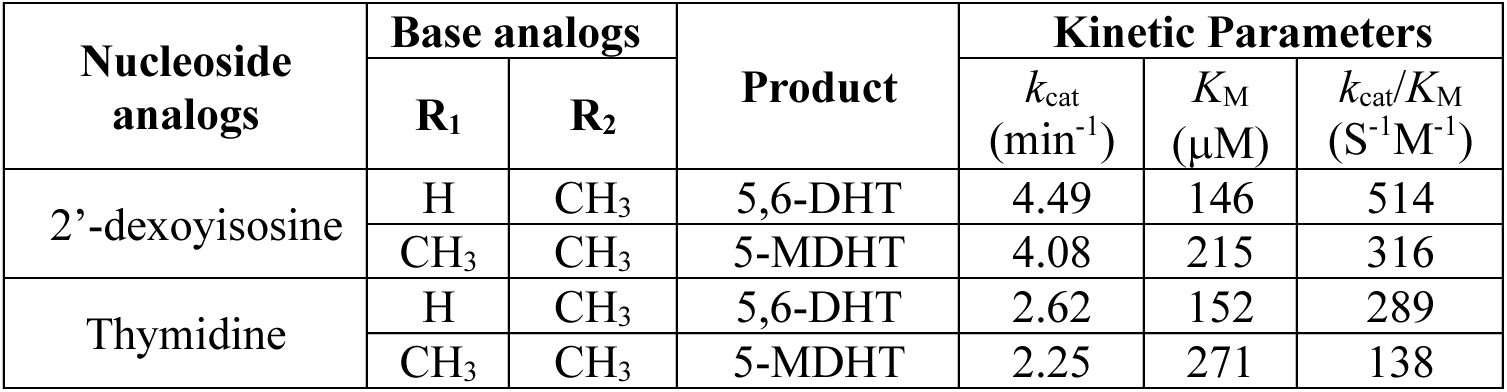
Relative Kinetics of hPNPase base-exchange catalysis reaction between uracil analogs and 2′-deoxyinosine or thymidine.

**Scheme 3:**
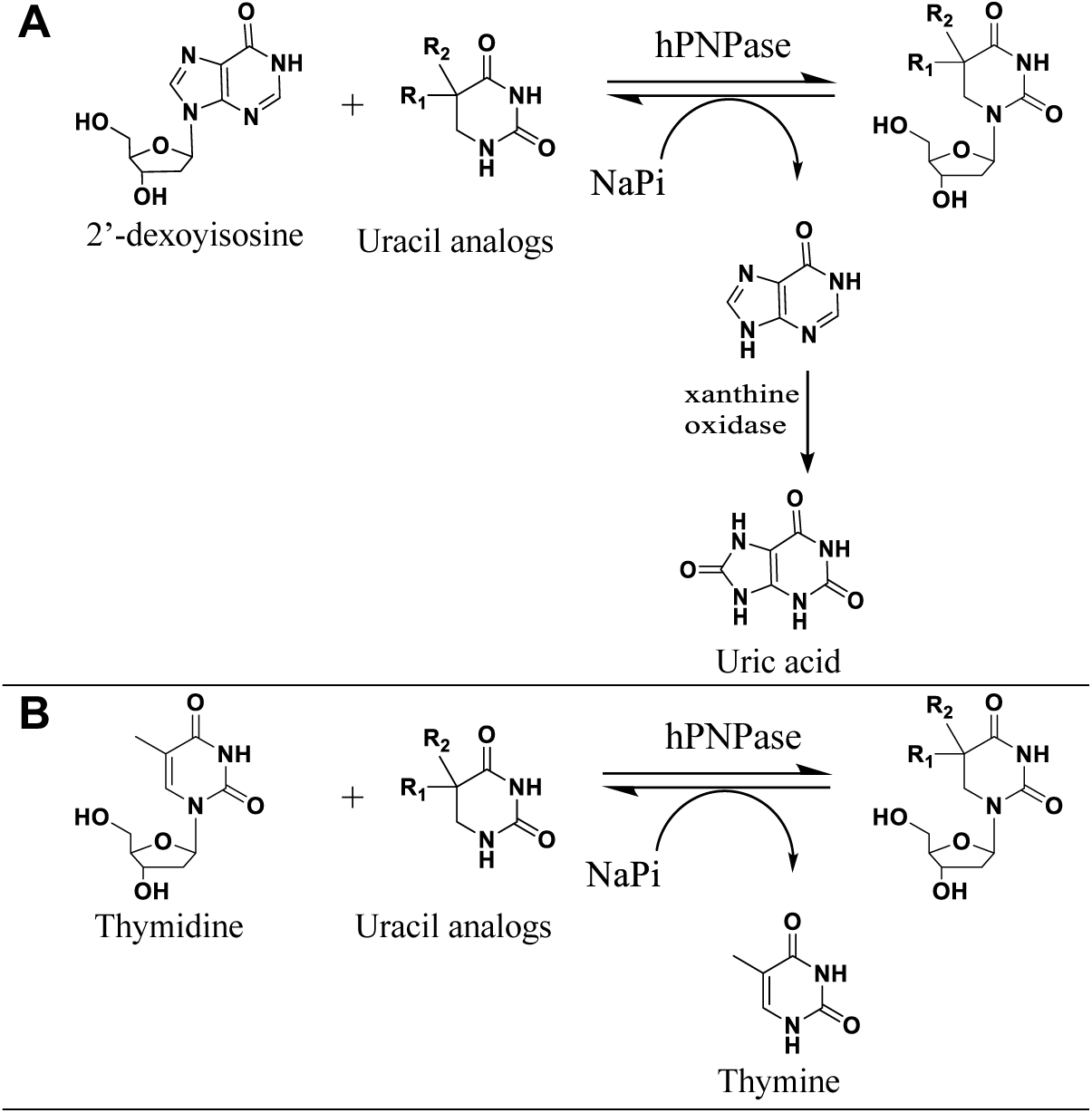
hPNPase base-exchange catalysis reaction between uracil analogs and 2’-dexoyisosine or thymidine.

Thymine base as a by-product will compete with the other uracil analogs (**7** and **8**) bases in the reaction, affecting the turnover number of the bio-catalyzed products by shifting the equilibrium reaction to the left. These calculated kinetic parameters of hPNPase base-exchange catalysis were used as a guide to scale up the production of 5,6-DHT and 5-MDHT and confirm their structures using NMR (**Figures S8-S13** of the Supporting Information).

#### Laboratory Scale-Up

The enzymatic assay with uracil analogs (**7** and **8**), hPNPase, and [2′-deoxyinosine/xanthine oxidase] or thymidine was scaled up to obtain (∼70 mg, 72% yield) of 5,6-DHT and (∼55 mg, 54% yield) of 5-MDHT. The ^1^H and ^13^C NMR spectra of the bio-catalyzed products suggested that hPNPase catalysis can be used to transfer 2’-deoxyribose from 2′-deoxyinosine or thymidine to the uracil substrates. This notion was supported by the chemical shifts of the protons and carbons bearing the C1′ of 2’-deoxyribose and the amino group N1 of the uracil base. The diagnostic NMR signal doublet of doublet observed at δ 6.29 for H1′ of the putative 5,6-DHT and 5-MDHT (**Figure 2D, 2E**, and **Figures S8-S13** of the Supporting Information) correspond to C1′ attached to an amino group of uracil base which similar to that for the standard thymidine (**Figure 2C**, and **Figure S18** of the Supporting Information). The ^13^C NMR results showed that the chemical shifts for C5 (δ 34.81) and C6 (δ 42,86) of 5,6-DHT and chemical shifts for C5 (δ 37.32) and C6 (δ 51.36) of 5-MDHT which further demonstrated the base exchange between the uracil analogs base and purine base (hypoxanthine) from 2′-deoxyinosine or pyrimidine base (thymine) from thymidine during the hPNPase catalysis. Moreover, LC-ESI-MS/MS monoisotopic-mass analysis validates the biocatalytic production of a correct molecular weight of [M-H]^-1^ at *m/z* 243.0972 for 5,6-DHT and *m/z* 257.1184 for 5-MDHT (**Figure S16** and **Figure S17** of the Supporting Information).

**Figure 2:**
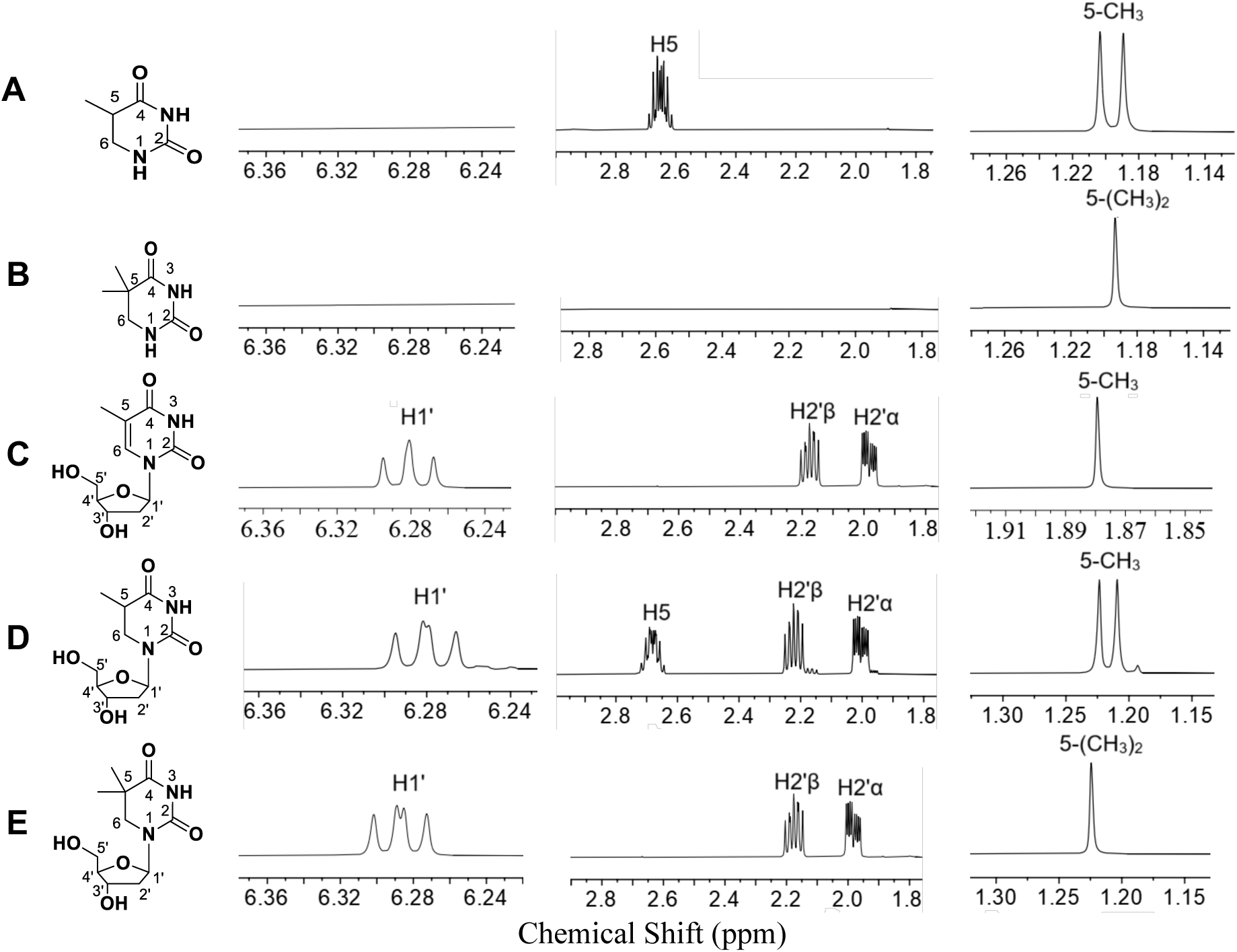
Partial 1H NMR (500 MHz, D_2_O) spectra of (A) 5,6-dihydro-5-methyluracil, (B) 5,5-dimethyl-1,3-diazinane-2,4-dione, (C) Thymidine, (D) 5,6-dihydrothymidine (5,6-DHT), and (E) 5-methyl-5,6-dihydrothymidine (5-MDHT).

### Activity and kinetics evaluation of SgvM^VAV^ and *Pa*HMT with 5,6-DHT as a substrate

#### Activity

The activity of SgvM^VAV^ and *Pa*HMT was tested by incubating 5,6-DHT as a substrate with SgvM^VAV^, *Pa*HMT, SAH, methyl iodide and with or without the addition of ZnCl_2_ to the assay reaction. LC-ESI-MS analysis was used to monitor bio-catalysis production. Selected ion *m/z* 257.11 was identified in the LC-ESI-MS profile of the assay reaction with ZnCl_2_ and assigned to the [M-H]^-1^ ion of the 5-MDHT (**Figure 3A**). At the same time, there is no production of 5-MDHT detected by the LC-ESI-MS for the assay reaction without ZnCl_2_ (**Figure 3B**), which indicates that ZnCl_2_ is required for the enzyme methyltransferase activity. Previous studies showed that Zn ions in the SgvM^VAV^ binding sites would stimulate a soft enolization in the α-keto compounds and, as a result, that will trigger a selective π-facial methyl transfer with the SAM compound and alkylate α-keto compounds.^33,34^

**Figure 3:**
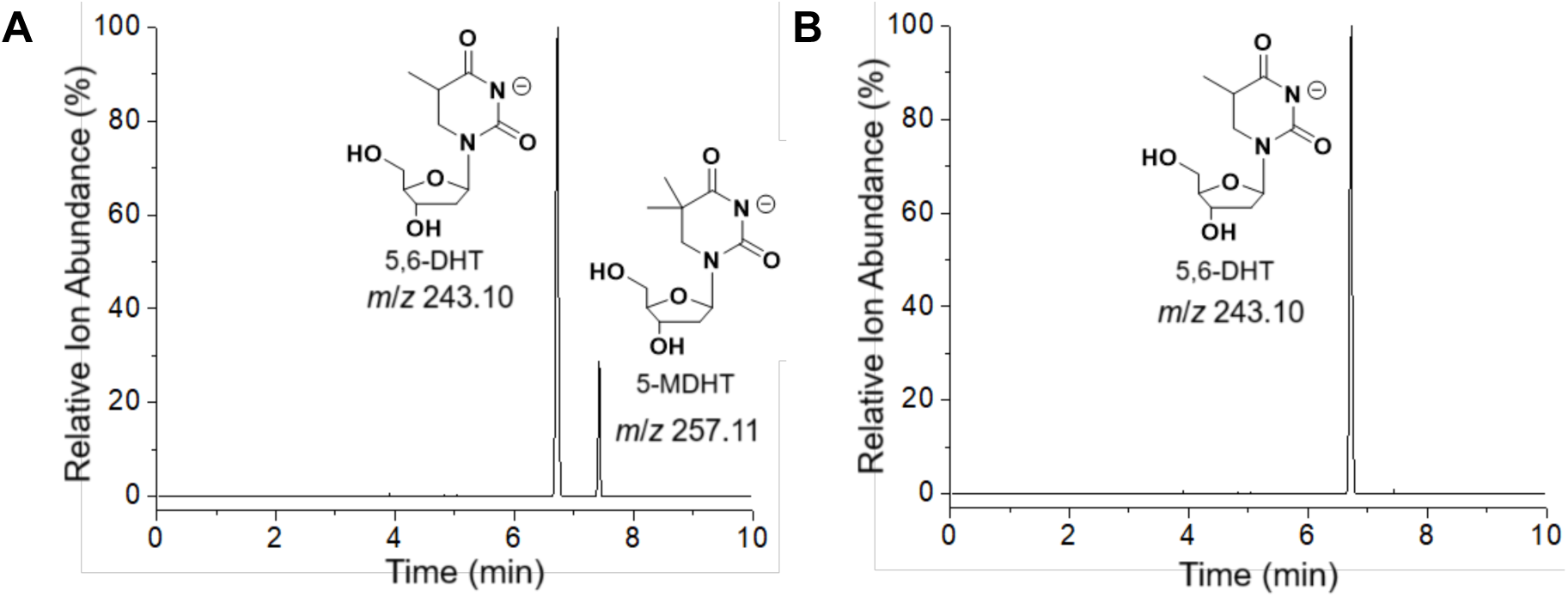
LC/ESI-MS (selected ion mode for *m/z* [M - H]^-^) of the biocatalytic conversion of 5,6-DHT to 5-MDHT made by SgvM^VAV^ and *Pa*HMT, (A) with ZnCl_2_ or (B) without ZnCl_2_.

#### Kinetics

Under steady-state conditions, the *K*_M_ and *k*_cat_ of the SgvM^VAV^ and *Pa*HMT catalysis were calculated by incubating SgvM^VAV^, *Pa*HMT, SAH, methyl iodide, and ZnCl_2_ with varying concentrations of 5,6-DHT (**Figure S7** of Supporting Information).

#### Laboratory Scale-Up

Based on the calculated kinetic parameters of the SgvM^VAV^ and *Pa*HMT catalysis, the biocatalytic production was scaled up to obtain (∼7 mg) of 5-MDHT. The ^1^H NMR results of the bio-catalyzed product (5-MDHT) showed a singlet observed at δ 1.23 ppm and integrated for 6 protons (**Figure 2E** and **Figure S14** of the Supporting Information) compared to that for the 5,6-DHT which is doublet at δ 1.22 ppm and integrated for 3 protons (**Figure 2D** and **Figure S8** of the Supporting Information). In addition, there is no chemical shift assigned for the H5 in the ^1^H NMR spectra of the 5-MDHT, which suggests that the C5 of the 5,6-DHT is alkylated by the SgvM^VAV^ catalysis (**Figure 2E** and **Figure S14** of the Supporting Information, see **Figure 2D** and **Figure S8** of the Supporting Information for comparison). Also, the ^13^C NMR result showed that the C5 chemical shift (δ 37.26) and C6 chemical shift (δ 51.35) for the 5-MDHT were shifted downfield (**Figure S11** of the Supporting Information) compared to those of 5,6-DHT which were shifted upfield C5 (δ 34.81) and C6 (δ 42.86) (**Figure S9** of the Supporting Information). These NMR results showed that SgvM^VAV^ and *Pa*HMT catalysis can be used to selectively transfer a methyl group to the C5 of the 5,6-DHT to create 5-MDHT. Furthermore, the LC-ESI-MS/MS monoisotopic-mass analysis verified a bio-catalyzed product of the correct molecular weight of [M-H]^-1^ at *m/z* 257.1046 for 5-MDHT (**Figure S17** of the Supporting Information) which supported the transfer of methyl group to the 5,6-DHT.

### Molecular dynamic simulation studies

#### Molecular modeling analysis of the hPNPase base-exchange catalysis reaction

A homology model of the hPNPase used in this study was constructed using the AMBER24 program. This model was based on the sequence homology and available crystal structure data of the human purine nucleoside phosphorylase (hPNPase) (PDB ID: 1RFG).^35^ The 5,5-dimethyl-5,6-dihydrouracil (**8**) with 2′-deoxyribose-1α-phosphate was docked in the active site of the hPNPase model using AutoDock Vina^36^, and UCSF Chimera^37^ to visualize and analyze all the binding poses. In this study, molecular dynamics simulations conducted a thermodynamics analysis on a series of conformations accessible to uracil analog (**8**) and the 2′-deoxyribose-1α-phosphate (αR1P) while docked in the hPNPase active site. Each substrate conformer’s intramolecular stability was calculated within the context of the proximate residues in the hPNPase active site. The MD simulations found several reasonably low energy-optimized conformational snapshots. These conformational snapshots aided in finding low-energy, catalytically competent structural conformations. Viewing a snapshot of these conformations shows that the Phe200, Met219, and Thr242 are interacting with the uracil analog (**8**) while His86 and Arg84 are interacting with the phosphate group of the 2′-deoxyribose-1α-phosphate (αR1P), which suggest that the substrates were likely positioned correctly as the catalytic residues aligned identically around the substrates as those in the structures of homologous enzyme from earlier studies (**Figure 4**).^38^

**Figure 4:**
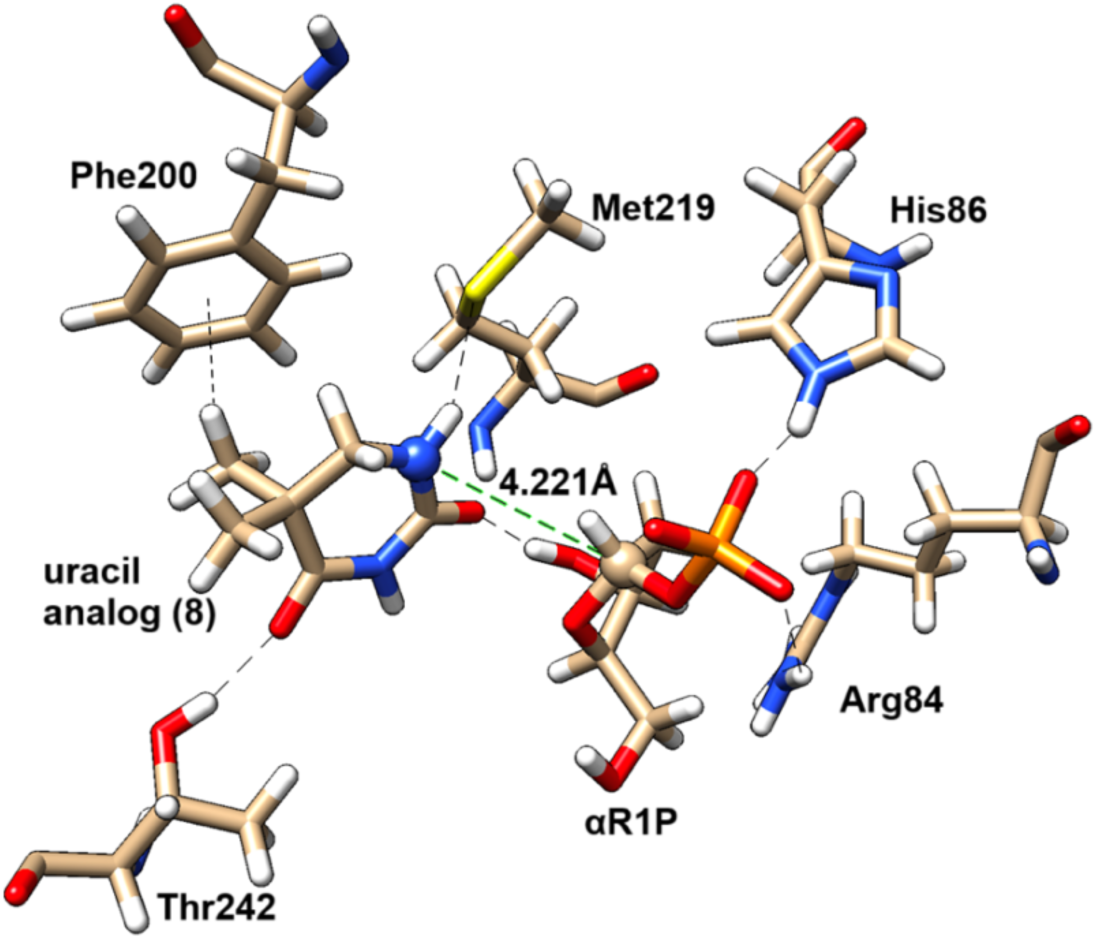
Structures of uracil analog (**8**) and the 2′-deoxyribose-1α-phosphate (αR1P) (the intermediate substrate in the hPNPase catalysis) within the hPNPase active site resulted from MD simulation. Here, the green dashed line shows the distance between the two reacting atoms (as noted in Scheme 2). Other black dashed lines show hydrogen bonds and methyl-π stacking interactions to stabilize the reactants in the hPNPase reaction pocket.

The human purine nucleoside phosphorylase (hPNPase) catalyzes the reversible cleavage of the glycosidic bond of 2′-deoxyinosine and thymidine, yielding the corresponding purine or pyrimidine bases and αR1P as products.^39–43^ The hPNPase can also catalyze the base-exchange reactions by N-glycosylated αR1P with the desired purine and pyrimidine to produce a range of unnatural nucleosides, including the 5-MDHT proposed in this study (**Scheme 4**).

**Scheme 4:**
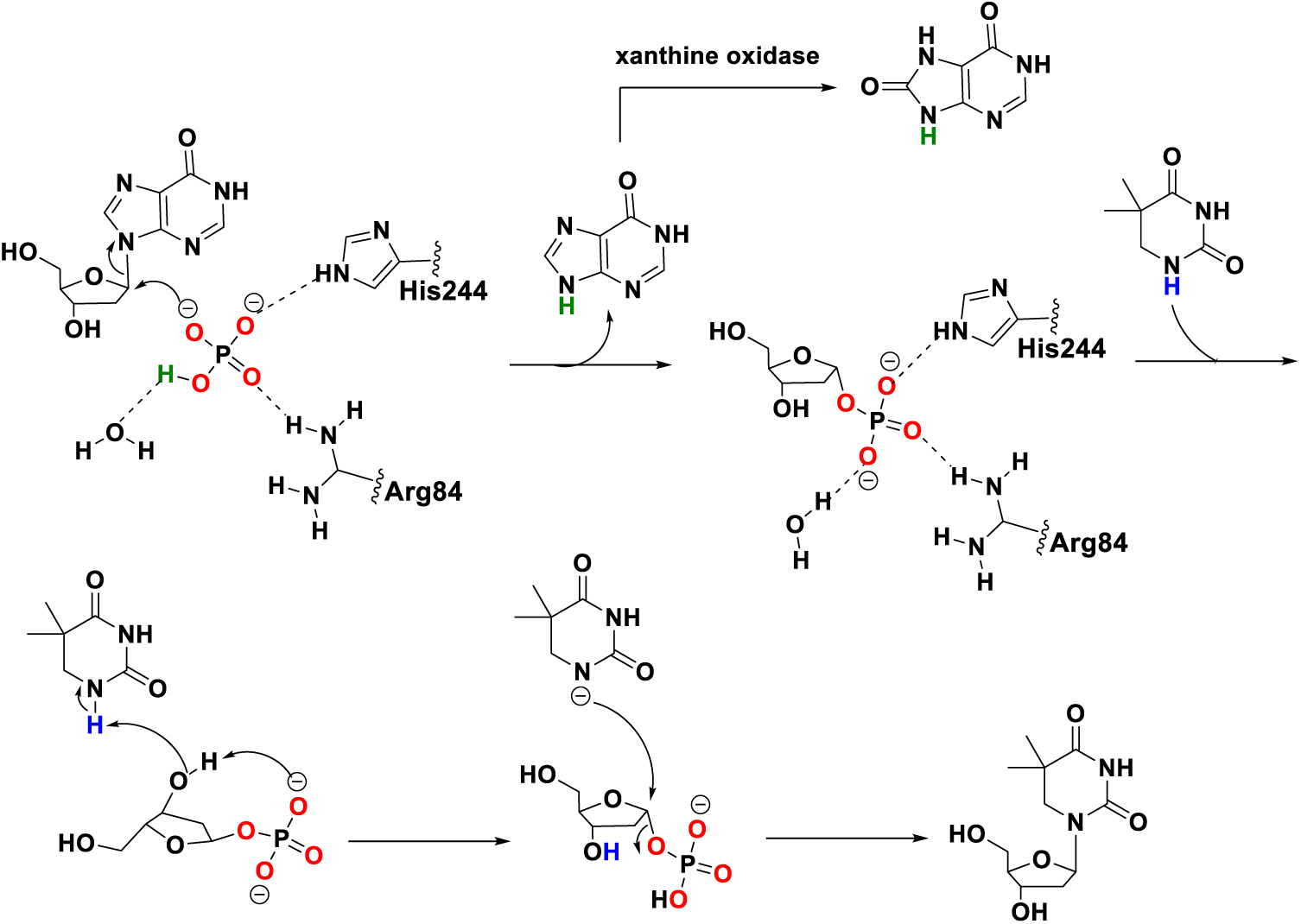
Proposed mechanism of hPNPase base-exchange catalysis between 2‵-deoxyinosine (**6**) and uracil analog (**8**) to produce 5-methyl-5,6dihydrothymidine (5-MDHT) (**5**).

To better understand the hPNPase base-exchange catalysis, we closely examined the uracil analog (**8**) bond to the hPNPase active site. The MD-simulation snapshot places the N1 of the uracil analog (**8**) at 4.2 Å from the C1¢ of the aR1P (**Figure 4**). A previous MD/extensive empirical valence bond (EVB) simulation study suggested that the purine base will be protonated/deprotonated through proton shuttle mechanisms.^44^ Our MD simulation shows that the N1-H of the uracil analog (**8**) is positioned at a suitable angle to prepare it for a possible proton shuttle mechanism with the C4-hydroxy and phosphate groups of the αR1P (**Figure 4**). After that, the N1 base of the uracil analog (**8**) will attack the C1′ of the αR1P through the S_N_2 mechanism fashioned (**Scheme 4**).

#### Molecular modeling analysis of the SgvM^VAV^ (SAM-methyltransferase) catalysis reaction

A homology model of SgvM^VAV^ used in this study was constructed using the AMBER24 program based on homology and available structure of SgvM^VAV^ (PDB ID: 8FTV). The Zn^2+^ ion, the Gaussian-optimized 5,6-DHT (**2**), and SAM were docked to the reaction site using AutoDock Vina and UCSF Chimera. MD simulations were carried out using the AMBER24 program, and several low-energy optimized conformational snapshots were generated. The intrinsic intermolecular and intramolecular stabilization energies of Zn^2+^ and the ligands were calculated within the context of the proximate residues in the enzyme active site. The low-energy snapshot suggested that the SgvM^VAV^ active site accommodates Zn^2+^ in an octahedral complex with the C4-carbonyl oxygen of 5,6-DHT, Asp245, His244, His297, and water (water hidden for better visual effect) (**Figure 5**). The Zn^2+^ ion acts as Lewis’s acid and can trigger soft enolization when it interacts with the C4-carbonyl oxygen of 5,6-DHT, which then facilitates electrophilic methylation (**Scheme 5**). The MD-simulation low-energy snapshot shows that the distance between the C5 position of 5,6-DHT and the SAM methyl group attached to sulfur atom is 4.5 Å, suggesting that SgvM^VAV^ could be able to transfer methyl group selectively to the C5 position of 5,6-DHT (**Figure 5**).

**Figure 5:**
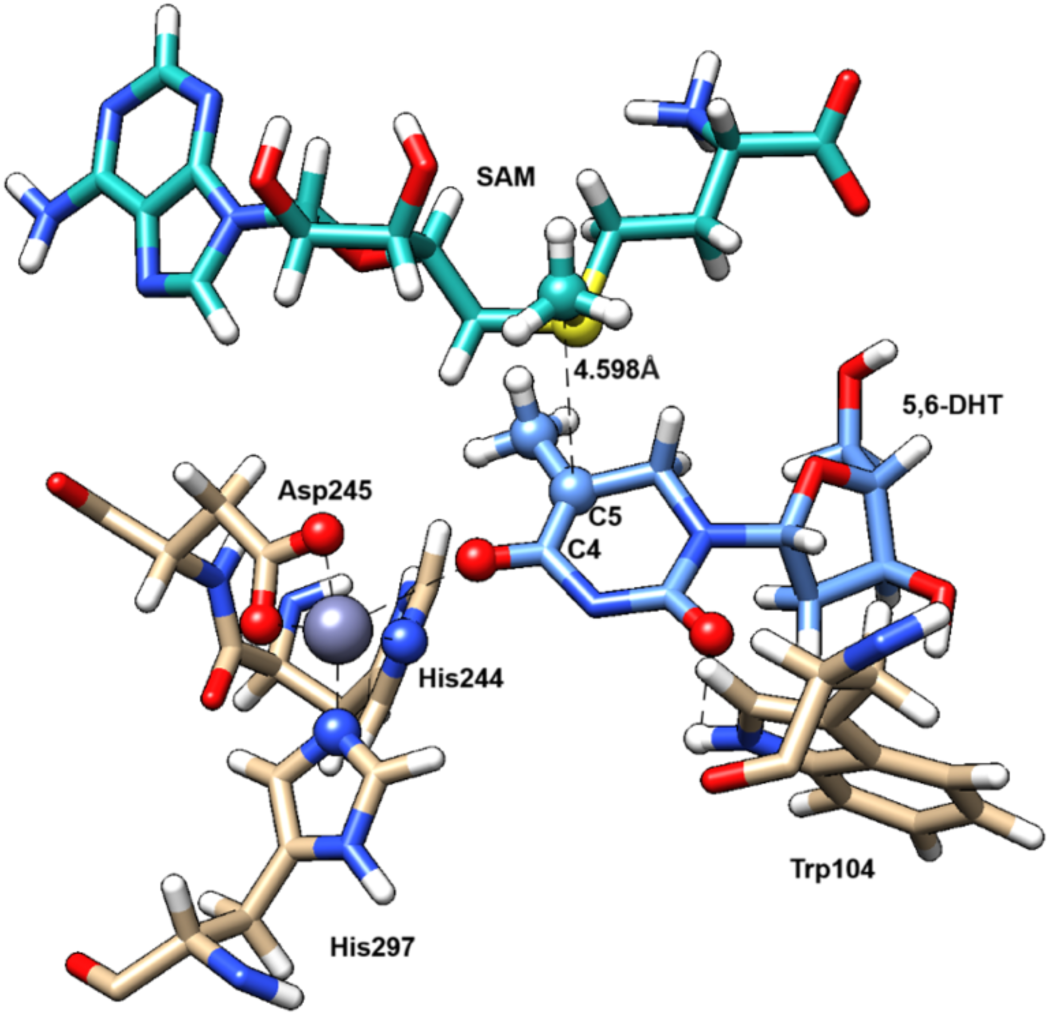
Structures of 5,6-dihydrothymidine (5,6-DHT) and S-Adenosyl methionine (SAM) within the SgvM^VAV^ active site resulted from MD simulation. The atoms estimated to form an octacoordinated complex with Zn^2+^ are drawn as spheres.

**Scheme 5:**
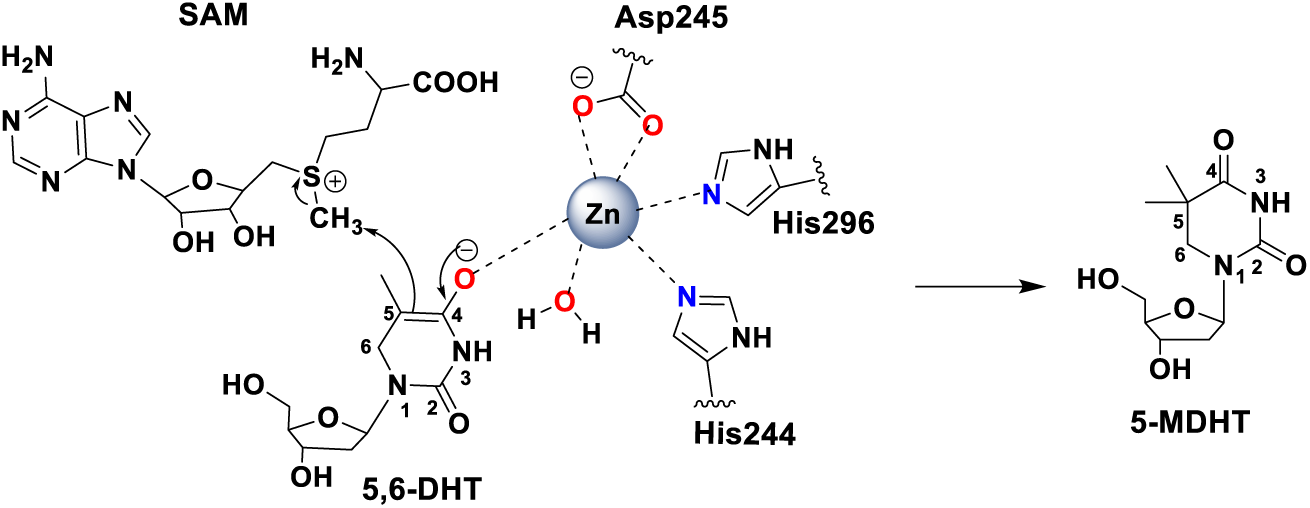
Proposed Zn^2+^-dependent alkylation mechanism of SgvM^VAV^ (SAM-methyltransferase) catalysis.

### Evaluating 5-MDHT as a CEST-MRI contrast agent

To evaluate whether 5-MDHT produced by biocatalysis is suitable as an MRI contrast agent, we compare two batches of 5-MDHT produced by two and single-step biocatalysis using CEST MRI. The two-step reaction was performed using the hPNPase base-exchange catalysis and then (SgvM^VAV^ and *Pa*HMT) methyltransferase catalysis with 5,6-DHT as an intermediate substrate (**Scheme 2A**). The single-step reaction was done using hPNPase base-exchange catalysis between uracil analogs and nucleoside substrates (**Scheme 2B**). In both experiments, the lyophilized 5-MDHT was dissolved in PBS (pH 7.4) for CEST MRI data collection. Both samples display the same typical peak at 5.3 ppm observable in the z-spectra (**Figure 6D-F**) and in the MTRasym (5.3 ppm) maps (**Figure 6C**). We also tested whether the catalyzed 5,6-dihydrothymidine (5,6-DHT) could be used as a CEST-MRI contrast agent. Notable, a peak at 5 ppm was observed in the z-spectra for the biocatalyzed 5,6-DHT (**Figure S19** of the Supporting Information). These results agree with the chemical synthesis results published by Bar-Shir *et al.* in 2013. The MTRasym map (**Figure 6C**) shows that the samples produce superior CEST contrast to thymidine due to the optimized exchange rate of imine proton. Therefore, the CEST MRI confirmed that biocatalysis can be used as an alternative method for the 5-MDHT chemical synthesis described in previous studies.

**Figure 6:**
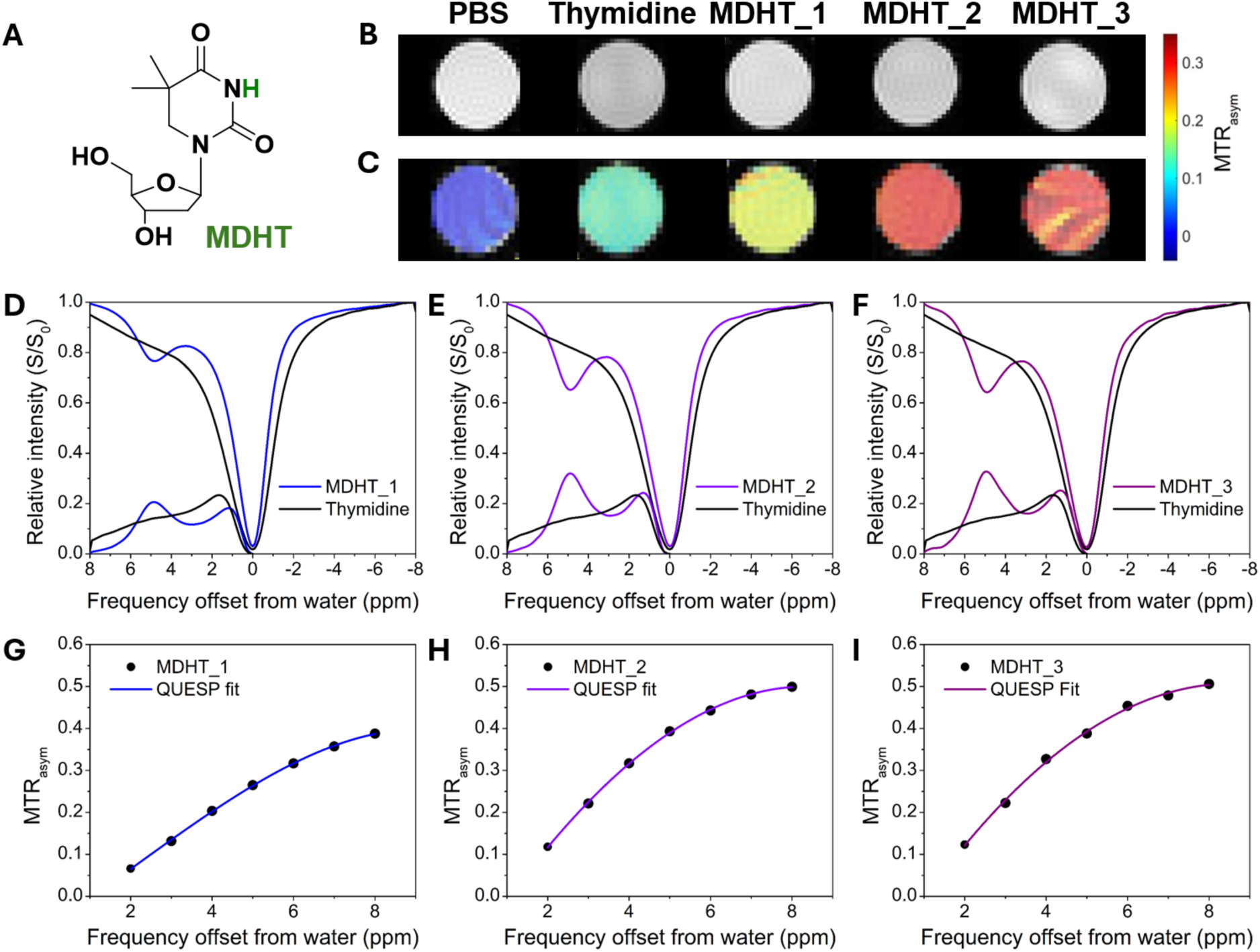
Evaluation of 5-MDHT as CEST MRI contrast agents. (A) The structure of 5-MDHT with the exchangeable imino proton is highlighted in green, (B) and (C) MTR_asym_ (5.3 ppm) maps, which show the contrast for the biocatalytic products of the two-step procedure (5-MDHT_1) and the single step-procedure (5-MDHT_2 and 5-MDHT_3). (D-F) CEST spectra and MTR_asym_ spectra of the three compounds. (G-I) MTR_asym_ (5.3 ppm) vs. saturation *B*_1_ (Hz), experimental (dots), and QUESP fitting (lines).

Genetically encoded reporters are widely used for monitoring molecular changes, with optical reporter genes being the most common. Over the years, various genes have been cloned from different organisms that emit light through bioluminescence or fluorescence.^45^ A new class of reporter genes encodes proteins that bind to radioisotopes or positron emitter probes, enabling quantitative imaging with appropriate radiolabeled probes.^46^ Therapeutic genes, such as the deoxynucleoside kinase family, can activate pro-drugs. Tjuvajev and Gambhi^47,48^ developed these into diagnostic genes for the nuclear imaging of HSV1-TK, a “theranostic” gene^49^, which has been clinically tested for imaging^50,51^ and therapy.^52,53^

MRI reporter genes present important advantages, as these enable the co-registration of gene expression information to be co-registered with anatomical and functional soft tissue data. They also enable minimally invasive, cell- and time-specific imaging of both eukaryotic and prokaryotic cells in vivo, achieving a spatial resolution of 50-100 µm.^54–56^ Initially developed for spectroscopy (MRS), these reporters also include proteins related to iron metabolism and storage^57–60^ and transgenic enzymes, which facilitate the conversion of compounds into paramagnetic contrast agents.^61,62^

Of the MRI reporters, genetically encoded reporters based on CEST-MRI have the added advantage that they can generate MRI contrast using bio-organic molecules, such as proteins, enzymes, and substrates, due to the biocompatible nature of the reporters. These provide greater flexibility in the design of the probe. Some CEST reporters rely on direct proton exchange with surrounding water protons^63–68^, while others are enzymes that catalyze substrates to produce the contrast. The enzymatic approach has two main advantages. The first is that it allows a large number of substrates for each enzyme, consequently increasing the number of contrast-generating protons. The second is that the substrates or imaging probes can be easily engineered. Simple modifications can shift the resonance frequency of the exchangeable protons away from that of the water protons, thereby reducing direct water saturation.

Additionally, the farther the frequency is, the lower the background generated from other metabolites in the cells, which improves the contrast-to-background ratio. The 5-MDHT exemplifies this, with the exchangeable imino proton resonating at a chemical shift of 5 ppm from the water proton. Another example is the substrate pyrrole-2’-deoxycytidine for the enzyme drosophila melanogaster 2′-deoxynucleoside kinase (Dm-dNK), where the chemical shift is even further at 6 ppm. The chemical shift can be pushed even farther away from the water proton resonance frequency by employing simulations and chemical modifications.^69^ However, due to its clinical relevance, we chose to use HSV1-TK in this study.

## Conclusions

Here, we demonstrated for the first time the synthesis of an MRI contrast agent using biocatalysis. Enzymes allow us to skip many traditional synthesis steps and reduce the generation of hazardous by-products and solvents. Moreover, optimizing enzyme activity through mutagenesis and protein modeling can improve the yield and specificity of the reaction. Thus, this approach represents a future avenue for the biosynthesis of other molecular imaging probes.

We developed an efficient method to produce the 5-MDHT imaging probe in a single-step reaction using hPNPase catalysis with high stereoselectivity and good yield. We examined the substrate’s scope and specificity of hPNPase catalysis. Our results showed that hPNPase could catalyze the base moiety of several nucleosides, such as thymidine and 2′-deoxyinosine, and transfer pyrimidine bases, such as uracil analogs (**7** and **8**) into the 1′-position of the nucleoside (2’-deoxyribose). Additionally, we used SgvM^VAV^ and *Pa*HMT catalysis as another method to produce the 5-MDHT by selectively installing a methyl group at the C5 position of the 5,6-DHT. The biocatalysis methods we proposed here to produce the 5-MDHT, a necessary CEST-MRI probe, could be used to develop many novel imaging probes for specific medical conditions. The biocatalysis method provides an alternative to chemical synthesis. This eco-friendly approach is faster, does not involve hazardous hydrogenation, and is easier to scale.

## Notes

### Competing Interest Statement

The authors have declared no competing interest.

### Funding

- NIH/NIBIB: R01-EB031008, R01-EB030565, R01-EB031936

## References

(1) Gambhir, S. S.; Barrio, J. R.; Phelps, M. E.; Iyer, M.; Namavari, M.; Satyamurthy, N.; Wu, L.; Green, L. A.; Bauer, E.; MacLaren, D. C.; Nguyen, K.; Berk, A. J.; Cherry, S. R.; Herschman, H. R. Imaging adenoviral-directed reporter gene expression in living animals with positron emission tomography. Proc. Natl. Acad. Sci. 1999, 96, 2333–2338.

(2) Tjuvajev, J. G.; Avril, N.; Oku, T.; Sasajima, T.; Miyagawa, T.; Joshi, R.; Safer, M.; Beattie, B.; DiResta, G.; Daghighian, F.; Augensen, F.; Koutcher, J.; Zweit, J.; Humm, J.; Larson, S. M.; Finn, R.; Blasberg, R. Imaging herpes virus thymidine kinase gene transfer and expression by positron emission tomography. Cancer Res. 1998, 58, 4333–4341.

(3) Gambhir, S. S.; Bauer, E.; Black, M. E.; Liang, Q.; Kokoris, M. S.; Barrio, J. R.; Iyer, M.; Namavari, M.; Phelps, M. E.; Herschman, H. R. A mutant herpes simplex virus type 1 thymidine kinase reporter gene shows improved sensitivity for imaging reporter gene expression with positron emission tomography. Proc. Natl. Acad. Sci. U.S.A. 2000, 97, 2785–2790.

(4) Ametamey, S. M.; Treyer, V.; Streffer, J.; Wyss, M. T.; Schmidt, M.; Blagoev, M.; Hintermann, S.; Auberson, Y.; Gasparini, F.; Fischer, U. C.; Buck, A. Human PET studies of metabotropic glutamate receptor subtype 5 with 11C-ABP688. J. Nucl. Med. 2007, 48, 247–252.

(5) Bar-Shir, A.; Alon, L.; Korrer, M. J.; Lim, H. S.; Yadav, N. N.; Kato, Y.; Pathak, A. P.; Bulte, J. W. M.; Gilad, A. A. Quantification and tracking of genetically engineered dendritic cells for studying immunotherapy. Magn. Reson. Med. 2018, 79, 1010–1019.

(6) Bar-Shir, A.; Liu, G.; Greenberg, M. M.; Bulte, J. W.; Gilad, A. A. Synthesis of a probe for monitoring HSV1-tk reporter gene expression using chemical exchange saturation transfer MRI. Nat. Protoc. 2013, 8, 2380–2391.

(7) Liu, G.; Liang, Y.; Bar-Shir, A.; Chan, K. W.; Galpoththawela, C. S.; Bernard, S. M.; Tse, T.; Yadav, N. N.; Walczak, P.; McMahon, M. T.; Bulte, J. W.; van Zijl, P. C.; Gilad, A. A. Monitoring enzyme activity using a diamagnetic chemical exchange saturation transfer magnetic resonance imaging contrast agent. J. Am. Chem. Soc. 2011, 133, 16326–16329.

(8) Ward, K. M.; Aletras, A. H.; Balaban, R. S. A new class of contrast agents for MRI based on proton chemical exchange dependent saturation transfer (CEST). Journal of magnetic resonance (San Diego, Calif. : 1997) 2000, 143, 79–87.

(9) Louie, A. Y.; Hüber, M. M.; Ahrens, E. T.; Rothbächer, U.; Moats, R.; Jacobs, R. E.; Fraser, S. E.; Meade, T. J. In vivo visualization of gene expression using magnetic resonance imaging. Nat. Biotechnol. 2000, 18, 321–325.

(10) Bar-Shir, A.; Liu, G.; Liang, Y.; Yadav, N. N.; McMahon, M. T.; Walczak, P.; Nimmagadda, S.; Pomper, M. G.; Tallman, K. A.; Greenberg, M. M.; van Zijl, P. C.; Bulte, J. W.; Gilad, A. A. Transforming thymidine into a magnetic resonance imaging probe for monitoring gene expression. J. Am. Chem. Soc. 2013, 135, 1617–1624.

(11) Yoo, B.; Pagel, M. D. A PARACEST MRI contrast agent to detect enzyme activity. J. Am. Chem. Soc. 2006, 128, 14032–14033.

(12) Yoo, B.; Raam, M. S.; Rosenblum, R. M.; Pagel, M. D. Enzyme-responsive PARACEST MRI contrast agents: a new biomedical imaging approach for studies of the proteasome. Contrast Media Mol. Imaging. 2007, 2, 189–198.

(13) Consolino, L.; Anemone, A.; Capozza, M.; Carella, A.; Irrera, P.; Corrado, A.; Dhakan, C.; Bracesco, M.; Longo, D. L. Non-invasive Investigation of Tumor Metabolism and Acidosis by MRI-CEST Imaging. Front. Oncol. 2020, 10, 161.

(14) Chen, Z.; Han, Z.; Liu, G. Repurposing Clinical Agents for Chemical Exchange Saturation Transfer Magnetic Resonance Imaging: Current Status and Future Perspectives. Pharmaceuticals (Basel, Switzerland) 2020, 14.

(15) Alon, L.; Kraitchman, D.; Schär, M.; Cortez, A.; Yadav, N.; Krimins, R.; Johnston, P.; McMahon, M.; van zijl, P.; Nimmagadda, S.; Pomper, M.; Bulte, J.; Gilad, A. Molecular Imaging of CXCL12 Promoter-driven HSV1-TK Reporter Gene Expression. Biotechnol. Bioprocess Eng. 2018, 23, 208–217.

(16) Chen, K.; Huang, X.; Kan, S. B. J.; Zhang, R. K.; Arnold, F. H. Enzymatic construction of highly strained carbocycles. Science. 2018, 360, 71–75.

(17) Brandenberg, O. F.; Fasan, R.; Arnold, F. H. Exploiting and engineering hemoproteins for abiological carbene and nitrene transfer reactions. Curr. Opin. Biotechnol. 2017, 47, 102–111.

(18) Al-Hilfi, A.; Walker, K. D. Biocatalysis of precursors to new-generation SB-T-taxanes effective against paclitaxel-resistant cancer cells. Arch. Biochem. Biophys. 2022, 719, 109165.

(19) Al-Hilfi, A.; Li, Z.; Merz Jr, K. M.; Nawarathne, I. N.; Walker, K. D. Biocatalytic and Regioselective Exchange of 2-O-Benzoyl for 2-O-(m-Substituted)Benzoyl Groups to Make Precursors of Next-Generation Paclitaxel Drugs. ChemCatChem. 2024, 16, e202400186.

(20) Al-Hilfi, A.; Li, Z.; Merz, K. M., Jr.; Walker, K. D. Mg2+-Ion Dependence Revealed for a BAHD 13-O-β-Aminoacyltransferase from Taxus Plants. JACS Au. 2024, 4, 4249–4262.

(21) Li, Y.; Li, S.; Thodey, K.; Trenchard, I.; Cravens, A.; Smolke, C. D. Complete biosynthesis of noscapine and halogenated alkaloids in yeast. Proc. Natl. Acad. Sci. U.S.A. 2018, 115, E3922–E3931.

(22) Teufel, R.; Kaysser, L.; Villaume, M. T.; Diethelm, S.; Carbullido, M. K.; Baran, P. S.; Moore, B. S. One-pot enzymatic synthesis of merochlorin A and B. Angew. Chem. Int. Ed. 2014, 53, 11019–11022.

(23) Frisch, M. J.; Trucks, G. W.; Schlegel, H. B.; Scuseria, G. E.; Robb, M. A.; Cheeseman, J. R.; Scalmani, G.; Barone, V.; Petersson, G. A.; Nakatsuji, H.; Li, X.; Caricato, M.; Marenich, A. V.; Bloino, J.; Janesko, B. G.; Gomperts, R.; Mennucci, B.; Hratchian, H. P.; Ortiz, J. V.; Izmaylov, A. F.; Sonnenberg, J. L.; Williams; Ding, F.; Lipparini, F.; Egidi, F.; Goings, J.; Peng, B.; Petrone, A.; Henderson, T.; Ranasinghe, D.; Zakrzewski, V. G.; Gao, J.; Rega, N.; Zheng, G.; Liang, W.; Hada, M.; Ehara, M.; Toyota, K.; Fukuda, R.; Hasegawa, J.; Ishida, M.; Nakajima, T.; Honda, Y.; Kitao, O.; Nakai, H.; Vreven, T.; Throssell, K.; Montgomery Jr., J. A.; Peralta, J. E.; Ogliaro, F.; Bearpark, M. J.; Heyd, J. J.; Brothers, E. N.; Kudin, K. N.; Staroverov, V. N.; Keith, T. A.; Kobayashi, R.; Normand, J.; Raghavachari, K.; Rendell, A. P.; Burant, J. C.; Iyengar, S. S.; Tomasi, J.; Cossi, M.; Millam, J. M.; Klene, M.; Adamo, C.; Cammi, R.; Ochterski, J. W.; Martin, R. L.; Morokuma, K.; Farkas, O.; Foresman, J. B.; Fox, D. J.: Gaussian 16 Rev. C.01. Wallingford, CT, 2016.

(24) Becke, A. D. Density-functional thermochemistry. III. The role of exact exchange. J. Chem. Phys. 1993, 98, 5648–5652.

(25) Lee, C.; Yang, W.; Parr, R. G. Development of the Colle-Salvetti correlation-energy formula into a functional of the electron density. Phys. Rev. B Condens. Matter. 1988, 37, 785–789.

(26) Tian, C.; Kasavajhala, K.; Belfon, K. A. A.; Raguette, L.; Huang, H.; Migues, A. N.; Bickel, J.; Wang, Y.; Pincay, J.; Wu, Q.; Simmerling, C. ff19SB: Amino-Acid-Specific Protein Backbone Parameters Trained against Quantum Mechanics Energy Surfaces in Solution. J. Chem. Theory Comput. 2020, 16, 528–552.

(27) Case, D. A.; Aktulga, H. M.; Belfon, K.; Cerutti, D. S.; Cisneros, G. A.; Cruzeiro, V. W. D.; Forouzesh, N.; Giese, T. J.; Götz, A. W.; Gohlke, H.; Izadi, S.; Kasavajhala, K.; Kaymak, M. C.; King, E.; Kurtzman, T.; Lee, T.-S.; Li, P.; Liu, J.; Luchko, T.; Luo, R.; Manathunga, M.; Machado, M. R.; Nguyen, H. M.; O’Hearn, K. A.; Onufriev, A. V.; Pan, F.; Pantano, S.; Qi, R.; Rahnamoun, A.; Risheh, A.; Schott-Verdugo, S.; Shajan, A.; Swails, J.; Wang, J.; Wei, H.; Wu, X.; Wu, Y.; Zhang, S.; Zhao, S.; Zhu, Q.; Cheatham, T. E., III; Roe, D. R.; Roitberg, A.; Simmerling, C.; York, D. M.; Nagan, M. C.; Merz, K. M., Jr. AmberTools. J. Chem. Inf. Model. 2023, 63, 6183–6191.

(28) Loncharich, R. J.; Brooks, B. R.; Pastor, R. W. Langevin dynamics of peptides: the frictional dependence of isomerization rates of N-acetylalanyl-N’-methylamide. Biopolymers. 1992, 32, 523–535.

(29) Berendsen, H. J. C.; Postma, J. P. M.; van Gunsteren, W. F.; DiNola, A.; Haak, J. R. Molecular dynamics with coupling to an external bath. J. Chem. Phys. 1984, 81, 3684–3690.

(30) Miyamoto, S.; Kollman, P. A. Settle: An analytical version of the SHAKE and RATTLE algorithm for rigid water models. J. Comput. Chem. 1992, 13.

(31) Dobritzsch, D.; Schneider, G.; Schnackerz, K. D.; Lindqvist, Y. Crystal structure of dihydropyrimidine dehydrogenase, a major determinant of the pharmacokinetics of the anti-cancer drug 5-fluorouracil. EMBO J. 2001, 20, 650–660-660.

(32) Smith, M. M.; Moran, G. R. The unusual chemical sequences of mammalian dihydropyrimidine dehydrogenase revealed by transient-state analysis. Meth. Enzymol. 2023, 685, 373–403.

(33) Ju, S.; Kuzelka, K. P.; Guo, R.; Krohn-Hansen, B.; Wu, J.; Nair, S. K.; Yang, Y. A biocatalytic platform for asymmetric alkylation of α-keto acids by mining and engineering of methyltransferases. Nat. Commun. 2023, 14, 5704.

(34) Sommer-Kamann, C.; Fries, A.; Mordhorst, S.; Andexer, J. N.; Müller, M. Asymmetric C-Alkylation by the S-Adenosylmethionine-Dependent Methyltransferase SgvM. Angew. Chem. 2017, 129, 4091–4094.

(35) Canduri, F.; Silva, R. G.; dos Santos, D. M.; Palma, M. S.; Basso, L. A.; Santos, D. S.; de Azevedo, W. F., Jr. Structure of human PNP complexed with ligands. Acta Crystallogr. B: Struct. Sci. Cryst. Eng. Mater. 2005, 61, 856–862.

(36) Eberhardt, J.; Santos-Martins, D.; Tillack, A. F.; Forli, S. AutoDock Vina 1.2.0: New Docking Methods, Expanded Force Field, and Python Bindings. J. Chem. Inf Model. 2021, 61, 3891–3898.

(37) Pettersen, E. F.; Goddard, T. D.; Huang, C. C.; Couch, G. S.; Greenblatt, D. M.; Meng, E. C.; Ferrin, T. E. UCSF Chimera--a visualization system for exploratory research and analysis. J. Comput. Chem. 2004, 25, 1605–1612.

(38) Pugmire, M. J.; Ealick, S. E. Structural analyses reveal two distinct families of nucleoside phosphorylases. Biochem. J. 2002, 361, 1–25.

(39) Komatsu, H.; Awano, H.; Ishibashi, H.; Oikawa, T.; Ikeda, I.; Araki, T. Chemo-enzymatic syntheses of natural and unnatural 2’-deoxynucleosides. Nucleic acids research. Supplement (2001) 2003, 101–102.

(40) Oslovsky, V. E.; Solyev, P. N.; Polyakov, K. M.; Alexeev, C. S.; Mikhailov, S. N. Chemoenzymatic synthesis of cytokinins from nucleosides: ribose as a blocking group. Org. Biomol. Chem. 2018, 16 12, 2156–2163.

(41) Zhang, W.; Turney, T.; Surjancev, I.; Serianni, A. S. Enzymatic synthesis of ribo- and 2′-deoxyribonucleosides from glycofuranosyl phosphates: An approach to facilitate isotopic labeling. Carbohydr. Res. 2017, 449, 125–133.

(42) Isaksen, G. V.; Åqvist, J.; Brandsdal, B. O. Thermodynamics of the Purine Nucleoside Phosphorylase Reaction Revealed by Computer Simulations. Biochem. 2017, 56, 306–312.

(43) Tebbe, J.; Bzowska, A.; Wielgus-Kutrowska, B.; Schröder, W.; Kazimierczuk, Z.; Shugar, D.; Saenger, W.; Koellner, G. Crystal structure of the purine nucleoside phosphorylase (PNP) from Cellulomonas sp. and its implication for the mechanism of trimeric PNPs. J. Mol. Biol. 1999, 294, 1239–1255.

(44) Isaksen, G. V.; Hopmann, K. H.; Åqvist, J.; Brandsdal, B. O. Computer Simulations Reveal Substrate Specificity of Glycosidic Bond Cleavage in Native and Mutant Human Purine Nucleoside Phosphorylase. Biochem. 2016, 55, 2153–2162.

(45) Shaner, N. C.; Campbell, R. E.; Steinbach, P. A.; Giepmans, B. N.; Palmer, A. E.; Tsien, R. Y. Improved monomeric red, orange and yellow fluorescent proteins derived from Discosoma sp. red fluorescent protein. Nat. Biotechnol. 2004, 22, 1567–1572.

(46) Serganova, I.; Ponomarev, V.; Blasberg, R. Human reporter genes: potential use in clinical studies. Nucl. Med. Biol. 2007, 34, 791–807.

(47) Tjuvajev, J. G.; Stockhammer, G.; Desai, R.; Uehara, H.; Watanabe, K.; Gansbacher, B.; Blasberg, R. G. Imaging the expression of transfected genes in vivo. Cancer Res. 1995, 55, 6126–6132.

(48) Gambhir, S. S.; Barrio, J. R.; Herschman, H. R.; Phelps, M. E. Imaging gene expression: principles and assays. J. Nucl. Cardiol. 1999, 6, 219–233.

(49) Song, Y.; Zou, J.; Castellanos, E. A.; Matsuura, N.; Ronald, J. A.; Shuhendler, A.; Weber, W. A.; Gilad, A. A.; Müller, C.; Witney, T. H.; Chen, X. Theranostics - a sure cure for cancer after 100 years? Theranostics. 2024, 14, 2464–2488.

(50) Jacobs, A. H.; Voges, J.; Reszka, R.; Lercher, M. J.; Gossmann, A.; Kracht, L.; Kaestle, C.; Wagner, R.; Wienhard, K.; Heiss, W.-D. Positron-emission tomography of vector-mediated gene expression in gene therapy for gliomas. The Lancet. 2001, 358, 727–729.

(51) Yaghoubi, S. S.; Jensen, M. C.; Satyamurthy, N.; Budhiraja, S.; Paik, D.; Czernin, J.; Gambhir, S. S. Noninvasive detection of therapeutic cytolytic T cells with 18F–FHBG PET in a patient with glioma. Nat. Clin. Pract. Oncol. 2009, 6, 53–58.

(52) Freytag, S. O.; Khil, M.; Stricker, H.; Peabody, J.; Menon, M.; DePeralta-Venturina, M.; Nafziger, D.; Pegg, J.; Paielli, D.; Brown, S.; Barton, K.; Lu, M.; Aguilar-Cordova, E.; Kim, J. H. Phase I study of replication-competent adenovirus-mediated double suicide gene therapy for the treatment of locally recurrent prostate cancer. Cancer Res. 2002, 62, 4968–4976.

(53) Freytag, S. O.; Stricker, H.; Peabody, J.; Pegg, J.; Paielli, D.; Movsas, B.; Barton, K. N.; Brown, S. L.; Lu, M.; Kim, J. H. Five-year follow-up of trial of replication-competent adenovirus-mediated suicide gene therapy for treatment of prostate cancer. Mol. Ther. 2007, 15, 636–642.

(54) Gilad, A. A.; Bar-Shir, A.; Bricco, A. R.; Mohanta, Z.; McMahon, M. T. Protein and peptide engineering for chemical exchange saturation transfer imaging in the age of synthetic biology. NMR in biomedicine 2023, 36, e4712.

(55) Duyn, J. H.; Koretsky, A. P. Novel frontiers in ultra-structural and molecular MRI of the brain. Curr. Opin. Neurol. 2011, 24, 386–393.

(56) Gilad, A. A.; Shapiro, M. G. Molecular Imaging in Synthetic Biology, and Synthetic Biology in Molecular Imaging. Mol. Imaging Biol. 2017, 19, 373–378.

(57) Genove, G.; DeMarco, U.; Xu, H.; Goins, W. F.; Ahrens, E. T. A new transgene reporter for in vivo magnetic resonance imaging. Nat. Med. 2005, 11, 450–454.

(58) Weissleder, R.; Moore, A.; Mahmood, U.; Bhorade, R.; Benveniste, H.; Chiocca, E. A.; Basilion, J. P. In vivo magnetic resonance imaging of transgene expression. Nat. Med. 2000, 6, 351–355.

(59) Cohen, B.; Ziv, K.; Plaks, V.; Israely, T.; Kalchenko, V.; Harmelin, A.; Benjamin, L. E.; Neeman, M. MRI detection of transcriptional regulation of gene expression in transgenic mice. Nat. Med. 2007, 13, 498–503.

(60) Matsumoto, Y.; Chen, R.; Anikeeva, P.; Jasanoff, A. Engineering intracellular biomineralization and biosensing by a magnetic protein. Nat. Commun. 2015, 6, 8721.

(61) Rallapalli, H.; McCall, E. C.; Koretsky, A. P. Genetic control of MRI contrast using the manganese transporter Zip14. Magn. Reson. Med. 2024, 92, 820–835.

(62) Bartelle, B. B.; Szulc, K. U.; Suero-Abreu, G. A.; Rodriguez, J. J.; Turnbull, D. H. Divalent metal transporter, DMT1: a novel MRI reporter protein. Magn. Reson. Med. 2013, 70, 842–850.

(63) Bar-Shir, A.; Bulte, J. W.; Gilad, A. A. Molecular engineering of nonmetallic biosensors for CEST MRI. ACS Chem. Biol. 2015, 10, 1160–1170.

(64) Gilad, A. A.; McMahon, M. T.; Walczak, P.; Winnard, P. T.; Raman, V.; van Laarhoven, H. W. M.; Skoglund, C. M.; Bulte, J. W. M.; van Zijl, P. C. M. Artificial reporter gene providing MRI contrast based on proton exchange. Nat. Biotechnol. 2007, 25, 217–219.

(65) Minn, I.; Bar-Shir, A.; Yarlagadda, K.; Bulte, J. W. M.; Fisher, P. B.; Wang, H.; Gilad, A. A.; Pomper, M. G. Tumor-specific expression and detection of a CEST reporter gene. Magn. Reson. Med. 2015, 74, 544–549.

(66) Fillion, A. J.; Bricco, A. R.; Lee, H. D.; Korenchan, D. E.; Farrar, C. T.; Gilad, A. A. Development of a synthetic biosensor for chemical exchange MRI utilizing in silico optimized peptides. NMR Biomed. 2023, 36, e5007.

(67) Farrar, C. T.; Buhrman, J. S.; Liu, G.; Kleijn, A.; Lamfers, M. L.; McMahon, M. T.; Gilad, A. A.; Fulci, G. Establishing the Lysine-rich Protein CEST Reporter Gene as a CEST MR Imaging Detector for Oncolytic Virotherapy. Radiol. 2015, 275, 746–754.

(68) Meier, S.; Gilad, A. A.; Brandon, J. A.; Qian, C.; Gao, E.; Abisambra, J. F.; Vandsburger, M. Non-invasive detection of adeno-associated viral gene transfer using a genetically encoded CEST-MRI reporter gene in the murine heart. Sci. Rep. 2018, 8, 4638.

(69) Yang, X.; Yadav, N. N.; Song, X.; Ray Banerjee, S.; Edelman, H.; Minn, I.; van Zijl, P. C. M.; Pomper, M. G.; McMahon, M. T. Tuning Phenols with Intra-Molecular Bond Shifted HYdrogens (IM-SHY) as diaCEST MRI Contrast Agents. Chem. Eur. J. 2014, 20, 15824–15832.

